# Ariadne: Synthetic Long Read Deconvolution Using Assembly Graphs

**DOI:** 10.1101/2021.05.09.443255

**Authors:** Lauren Mak, Dmitry Meleshko, David C. Danko, Waris N. Barakzai, Salil Maharjan, Natan Belchikov, Iman Hajirasouliha

## Abstract

Synthetic Long Read (SLR) sequencing techniques such as UST’s TELL-Seq, and Loop Genomics’ LoopSeq combine 3^′^ barcoding with standard short-read sequencing to expand the range of linkage resolution from hundreds to tens of thousands of base-pairs. However, the lack of a 1:1 correspondence between a long fragment and a 3^′^ unique molecular identifier (UMI) confounds the assignment of linkage between short-reads. We introduce Ariadne, a novel assembly graph-based SLR deconvolution algorithm, that can be used to extract single-species read-clouds from SLR datasets to improve the taxonomic classification and *de novo* assembly of complex populations, such as metagenomes.

## 1 Background

Next generation sequencing technologies underpin the large-scale genetic analyses of complex mixtures of isolates, such as microbiomes. However, the simultaneous reconstruction of multiple distinct and discrete genomes, especially in communities where the number of species is unknown, is much more computationally demanding than assembling a single isolate genome.

Though standard Illumina short-read sequencing is the most popular platform for metagenome-wide characterizations due to its sequencing depth to cost ratio, the length of Illumina short-reads limits the linkage information that can be extracted from sequencing libraries. To address this, researchers are increasingly using nucleotide barcode-based, chromatin conformation-based, or long-read sequencing technologies to approximate single-cell resolution from complex mixtures of microorganisms. Long-read sequencing technologies, such the range of options from Oxford Nanopore Technologies, are capable of resolving a limited number of high-quality circularized draft genomes from metagenomic sequencing data [1]. However, highquality assemblies of long reads are reliant on large amounts of input DNA to adequately span all of the species in a metagenomics sample, which may not be available depending on the size of the individual genomes and the alpha diversity [2]. When sequencing depth is low, the assembly graph contains many ambiguous junctions comprised of sequence material from multiple genetically similar species and/or strains [1]. Low sequencing coverage means that some edges in the *de novo* assembly de Bruijn graph are interpreted as sequencing errors, and mistakenly removed.

### 1.1 Overview of Synthetic Long Read (SLR) Technology

Alternative approaches that rely on short-read sequencing while still generating long-range linkage information include Hi-C and synthetic long-read (SLR) technologies [3]. While similar ambiguous junctions are inherent in short-read assemblies due to limitations in sequencing insert size, *de novo* assemblies generated by short-read and synthetic long read (SLR) are inherently larger and more comprehensive representations of the sample composition due to the relative volume of genomic coverage. Since SLR library preparations require less input material than long reads, it is much more suitable for extracting sequencing information from low-volume samples [1, 4, 5, 6].

SLR technologies, starting with Illumina TruSeq Synthetic Long reads, [7], associate short reads originating from the same extracted genomic fragment with some type of nucleotide-based unique molecular identifier (UMI). UMIs are colloquially referred to as ‘barcodes’ in the context of SLR sequencing. Briefly, input (meta)genomic DNA is sheared into long fragments of 5-100 kbp. After shearing, a UMI (usually 16 - 20 base-pairs long) is ligated to short-reads from the fragments such that short-reads from the same fragment share the same UMI. Linked-read UMIs are unrelated to the 5’ UMI used for sample multiplexing. We refer to the set of reads that share a UMI as a read cloud. Finally, the short-reads are sequenced using standard sequencing technologies (e.g. Illumina HiSeq). SLRs offer additional long-range information over standard shortreads. Reads with matching UMIs are more likely to have emerged from the same long fragment of DNA than two randomly sampled reads, which extends the relative positional information encoded in the read’s short 50 - 250 bp sequence past the standard limitations of a short-read insert, which are typically several hundreds of base-pairs. There is a slight coverage tradeoff due to the size of the UMIs and associated library preparation costs relative to standard short-read sequencing.

SLR techniques are differentiated by the biochemical mechanism that associates UMIs with genomic fragments or short reads. 10x linked-read sequencing uses oil-based droplets to encapsulate a few genomic fragments and a UMI, which are subsequently splintered into short reads which are amplified with the UMI [8]. While 10x Genomics’ linked reads has been discontinued after nearly half a decade of widespread usage, a variety of other UMI-based SLR methods such as TELL-Seq, LoopSeq, and BGI’s Long Fragment Reads (stLFR) have been commercialized recently with the promise of (near-)single-molecule resolution along with simplified library preparation procedures and compatibility with standard Illumina sequencing machines. TELL-Seq uses proprietary TELL beads, which both capture genomic fragments and use transposon-based reactions to insert the UMI sequence throughout the fragment [9]. TELL fragments are approximately 20 - 40 kbp long [9], which are similar in length to the (on average) 10 kbp fragments generated by 10x [10]. LoopSeq similarly uses an intramolecular enzyme-based distribution method, but does not use beads to partition genomic fragments at all [11]. One major advantage of both TELL-Seq and LoopSeq over 10x and BGI’s Long Fragment Reads is their single-tube reaction chemistry, and compatibility with standard Illumina sequencing machines. 10x library preparation depended on costly instruments for droplet management [8], and BGI’s Long Fragment Reads are incompatible with Illumina sequencing. For detailed explanations of each method’s capabilities and library preparation procedures, we direct the reader to [8, 9, 11, 12]. In this study, we also provide a first head-to-head comparison of multiple SLR technologies to profile their strengths and weaknesses on metagenomics datasets of similar complexity.

The central drawback of SLR technologies is that the UMI-based linkage information must be efficiently interpreted from the sequencing data to simulate long read resolution and increase the average contiguity (i.e.: NA50) of *de novo* assembly. The need for novel algorithms to leverage this information has been partially met by the proliferation of SLR-based assemblers, such as Athena, Supernova, and cloudSPAdes [13, 14, 15]. However, additional novel algorithmic efforts are needed to achieve the desired contiguity and reduce the number of observed assembly errors [15]. Many taxonomic lineages identified in large-scale studies are not represented in reference sequence databases, and are not associated with isolated cultures [16, 17, 18]. Metagenomic analyses that rely on existing reference genomes, such as read-based taxonomic classification, will inherently bias the genomic reconstruction of a mixed population, generally towards the most common and already well-characterized species within the sample [19]. Thus, reference-based analysis is not ideal as a first step for samples with large variations in species compositions or samples from substrates that are poorly represented in reference databases, such as under-studied or extreme environments [20, 18].

### 1.2 Applications of SLR Sequencing in Metagenomics

SLR sequencing has been shown to resolve species compositions of metagenomics samples in both 16S-based and shotgun whole-genome-based analyses. Because of its low UMI multiplicity, LoopSeq was able to identify multiple copies of the 16S rRNA gene as belonging to a single strain [21, 22]. When paired with additional sources of information, such as targeted amplification or longitudinal sequencing, SLRs are capable of resolving metagenomics to the strain level. In conjunction with high-throughput qPCR, LoopSeq has been used to associate antimicrobial resistance genes with specific species in environmental samples [23]. The long-read-like linkage information encoded in read clouds has been used to track single nucleotide variants in the human gut microbiome longitudinally, demonstrating that prioritizing depth of coverage over strict read length can be an optimal analysis strategy especially when the number of strains/haplotypes is an unknown variable [10]. In another study, SLRs have also been used to identify the presence of structural variation on bacterial chromosomes [24].

### 1.3 Challenges Posed by SLR Sequencing

Despite the additional linkage information offered by SLR sequencing, there are new computational challenges involved in applying barcoded reads to *de novo* assembly. Because metagenomic samples are intrinsically multiplexed samples of multiple species, longrange linkage information is confounded by the multiplicity of fragments assigned to 3^′^ UMIs. Existing systems employ on the order of 10^6-7^ 3^′^ UMIs [25]. In previous studies with the 10x Genomics system, it was observed that there were 2–20 long fragments of DNA per 3^′^ UMI [26]. The larger the barcoded read cloud, the more likely that reads tended to originate from multiple fragments. Our analyses suggest that at least 97% of read clouds are composed of *≥*2 fragments, with the exception of a LoopSeq dataset. In the absence of additional information about the sample, it is difficult to distinguish the genomic origin of a random assortment of reads with the same UMI. Furthermore, each fragment of DNA is only fractionally covered by reads (typically 10-20%). Because overall coverage is reduced, SLRs provide long-range information at the expense of short-range blocks of contiguous sequence.

### 1.4 The Barcode Deconvolution Problem

The barcode deconvolution problem, previously described in [27, 26], is defined as the assignment of each read with a given 3^′^ UMI to a subgroup such that every read in the subgroup originates from the same fragment or contiguous genomic segment. A solution to the barcode deconvolution problem for a set of read clouds would be a map from each read cloud to a function which solves the UMI deconvolution problem for that read cloud. Reads with the same 3^′^ UMI that are highly likely to have originated from the same metagenomic fragment are more likely to co-occur in one another’s sequence space within the assembly graph than reads from different fragments.

Though the multiplicity of genomic fragments to 3^′^ UMIs has been problematic since the inception of SLR sequencing, the UMI deconvolution problem has only been addressed recently by two computational tools: EMA [27] and Minerva [26]. The EMA approach augments read alignment for barcoded reads based on alignment to a reference sequence. In the process of generating probabilistic alignments, reads from a single read cloud are sub-grouped into sets of reads that map close to each other. EMA is particularly applicable for highly repetitive regions where a barcoded read can align to multiple locations within the genome [27]. However, the EMA approach relies on the user to supply reference sequence(s) as *a priori* information about the input sample. We also demonstrate that EMA (as of the date of publication) is unable to recognize > 99.9% of SLR UMIs, even with UMI whitelists tailored for each dataset. Additionally, [28] has proposed an extension of the EMA approach that makes a graph of UMIs using the degree of read connectivity between UMIs as edges to deconvolve read clouds. While the species composition of popular sampling sites such as the human gut are well-characterized, such reference-based methods are not designed for microbial samples from under-studied environments such as urban landscapes [20]. Minerva does not require a reference genome, using instead *k*-mer similarities between read clouds to approximately solve the UMI deconvolution problem for metagenomic samples [26]. However, Minerva is memory-inefficient and requires extensive parameter optimization to deconvolve all of the read clouds in a dataset.

In this paper, we present Ariadne as an advanced approach to tackle UMI deconvolution. Instead of using read alignments to reference sequences, which are unknown for poorly characterized environmental microbial samples [20], or relying on computationally expensive string-based graphs, Ariadne leverages the linkage information encoded in the full de Bruijn-based assembly graph generated by a *de novo* assembly tool such as cloudSPAdes [15] to generate up to 37.5-fold more read clouds containing only reads from a single fragment, improve the summed NA50 by up to 500 kbp, and maintain a proportional rate of misassembly relative to *de novo* assembly without prior UMI deconvolution. Searching through the pre-made assembly graph for reads in the neighboring sequence space makes the search for co-occurring reads computationally tractable, generalizable, and scalable to large datasets.

## 2 Results

### 2.1 Benchmarking Datasets

We used real data sets from four microbial mock communities which we refer as MOCK5 10x, MOCK5 LoopSeq, MOCK20 10x, MOCK20 TELL-Seq throughout this manuscript. See Table 1 for an overview of the mock microbiome datasets and Table 2 for the relative species abundances. Two of the datasets, MOCK5 LoopSeq and MOCK20 TELL-Seq, are analysed for the first time in this work. Note that while both comprised of 5 species, the MOCK5 10x and MOCK5 LoopSeq datasets share only one species—*Escherichia coli* —and thus cannot be considered as a comparison of 10x and LoopSeq technology. The original MOCK20 10x dataset is approximately 330 million reads, but for efficiency a random subset of 100 million reads was used in the subsequent analyses. Similarly, the original MOCK20 TELL-Seq dataset is approximately 210 million reads large, but was also subsetted to 100 million reads. Reference genome sequences can be downloaded [here](https://github.com/lauren-mak/ariadne/blob/ariadne_1.0/reference_seqs.tar.gz).

**Table 1.**
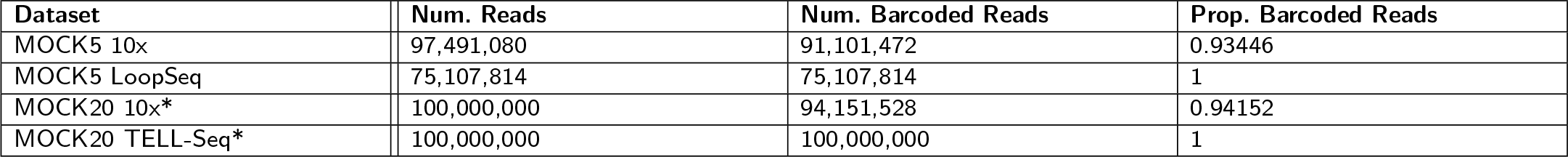
Overview of mock microbiome linked-read datasets. * indicates that MOCK20 10x and TELL-Seq were generated from the same mock microbiome community product.

| Dataset | Num. Reads | Num. Barcoded Reads | Prop. Barcoded Reads |
| --- | --- | --- | --- |
| MOCK5 10x | 97,491,080 | 91,101,472 | 0.93446 |
| MOCK5 LoopSeq | 75,107,814 | 75,107,814 | 1 |
| MOCK20 10x* | 100,000,000 | 94,151,528 | 0.94152 |
| MOCK20 TELL-Seq* | 100,000,000 | 100,000,000 | 1 |

**Table 2.** Relative species abundance in each mock microbiome community as calculated by the tool CoverM [41]. Abundance based on read coverage of reference genome sequence adjusted for genome size.

| Dataset | Species | Relative Abundance |
| --- | --- | --- |
| MOCK5 10x | Enterobacter cloacae | 7.129894635 |
|  | Pseudomonas fluorescens | 11.63274034 |
|  | Micrococcus luteus | 16.54618068 |
|  | Escherichia coli | 22.04082093 |
|  | Staphylococcus epidermidis | 42.6503661 |
| MOCK5 LoopSeq | Pseudomonas aeruginosa | 2.2308808 |
|  | Streptococcus mutans | 5.312954304 |
|  | Porphyromonas gingivalis | 26.40226344 |
|  | Escherichia coli | 28.9818004 |
|  | Rhodobacter sphaeroides | 37.07210106 |
| MOCK20 10x and TELL-Seq | Schaalia odontolytica | 0.005069583 |
|  | Bifidobacterium adolescentis | 0.005194844 |
|  | Deinococcus radiodurans | 0.014251377 |
|  | Enterococcus faecalis | 0.015894382 |
|  | Bacteroides vulgatus | 0.027645357 |
|  | Cutibacterium acnes | 0.184333912 |
|  | Lactobacillus gasseri | 0.189651678 |
|  | Neisseria meningitidis | 0.229268868 |
|  | Helicobacter pylori | 0.241180209 |
|  | Acinetobacter baumannii | 0.250956306 |
|  | Bacillus cereus | 1.214779563 |
|  | Clostridium beijerinckii | 1.220119982 |
|  | Pseudomonas aeruginosa | 1.366135524 |
|  | Staphylococcus aureus | 1.816064537 |
|  | Streptococcus agalactiae | 1.914207527 |
|  | Rhodobacter sphaeroides | 13.2715074 |
|  | Escherichia coli | 16.14207583 |
|  | Staphylococcus epidermidis | 19.55632719 |
|  | Porphyromonas gingivalis | 21.07923273 |
|  | Streptococcus mutans | 21.25610476 |

### 2.2 Gold-Standard (‘Reference’) Cloud Deconvolution

The use of mock communities allowed us to infer the genomic fragment of origin of the barcoded reads. These gold-standard fragment assignments served as a benchmark to compare read clouds and assemblies with and without deconvolution. We mapped the reads to the reference sequences of the species that were known to form the mock communities using Bowtie2 [29] (version 2.3.4.1). Using the read mapping positions along the reference genome, we further subsetted the reads into gold-standard read clouds such that the left- and right-most starting positions of reads in the same cloud were no further than 200 kbp apart, in case multiple fragments from the same genome were present in the same read cloud. We termed this method ‘reference deconvolution’ as it represents the database/reference-sequence-based inference of the genomic fragments that originated reads tagged with the same 3^′^ UMI.

### 2.3 Ariadne Generates a Large Number of High-Quality Enhanced Read Clouds

Ariadne generated enhanced read clouds, or subgroups of original read clouds, that largely corresponded to individual fragments of DNA. We measured the quality of each enhanced read cloud using two metrics: Shannon entropy index H = Σ*p_i_* log *p_i_* and purity 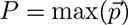 where *p_i_* indicates the proportion of reads in an (enhanced) read cloud that originates from the same most prevalent 200 kbp region in a single reference sequence. It should be noted that the reference-based clouds are presented here as a high-water comparison only. If the species composition of the original sample were already known, barcode deconvolution is not applicable. Prior to these quality checks, we excluded reads from standard and enhanced read groups of size 2 or smaller (i.e. consisting of a single read-pair or smaller), which are trivially pure.

Without deconvolution, there is a large spread of P at each read cloud size, and nearly 100% of the reads are in mixed-origin read clouds, or clouds that are comprised of reads that have likely originated from different SLR fragments (Table 3). Larger clouds are more likely to contain reads of multiple species origins and thus lower purity. The exception to this trend is the MOCK5 LoopSeq dataset (Figure 1 top row). Since the goal of linked reads is to approximate the linkage range of long fragments, having mixed-origin clouds as a result of fragment-to-UMI multiplicity confounds downstream applications such as taxonomic classification. Ariadne deconvolution reduces the proportion of multi-origin read clouds by at least 2-fold in the MOCK5 LoopSeq dataset, up to 7.5-fold in the MOCK20 10x dataset (Table 3 column 7). Since Ariadne deconvolution generally decreases this trend (except in the case of LoopSeq), it is unlikely that our results have been inflated by a large number of small and trivially pure clouds. Furthermore, reference-based deconvolution generates a similar number of clouds of a similar size to Ariadne, indicating that search-distance based subgrouping models genomic fragment boundaries sufficiently. The exception to this trend is the MOCK20 10x dataset, where Ariadne generated twice the number of read clouds that are slightly less than half the size of clouds in the reference-deconvolved dataset (Table 3). In general, Ariadne produces enhanced read clouds that are smaller than reference-deconvolved read clouds, indicating that on average, reads that should be included in the cloud are ‘missing’ and that the search-distance-based method does not exhaustively cluster all reads from the same fragment.

**Table 3.** Read cloud summary statistics. For Ariadne deconvolution, we used a search distance of 5 kbp. The 3-5^th^ columns contain the average and standard deviations of read cloud statistics. Prop. Under, Comp., and Over refer to the proportion of total original or deconvolved read clouds that were over- or under-deconvolved, or completely and exactly comprised of all of the reads from a single inferred genomic fragment.

| Dataset | Deconv. Method | Avg. Purity | Avg. Entropy | Avg. Size | Num. Clouds | Prop. Under | Prop. Comp. | Prop. Over |
| --- | --- | --- | --- | --- | --- | --- | --- | --- |
| MOCK5 10x | None | 0.53±0.16 | 1.61±0.52 | 63.51 ±51.64 | 1 425 430 | 0.98 | 0.02 | 0 |
| MOCK5 10x | Reference | 0.99±0.05 | 0.03±0.14 | 18.47 ±19.26 | 4 733 136 | 0.06 | 0.91 | 0.03 |
| MOCK5 10x | Ariadne | 0.89±0.22 | 0.37±0.7 | 12.18 ±15.26 | 4 753 473 | 0.25 | 0.04 | 0.71 |
| MOCK5 LoopSeq | None | 0.93±0.11 | 0.31±0.31 | 269.18 ±356.67 | 276 665 | 0.68 | 0.32 | 0 |
| MOCK5 LoopSeq | Reference | 0.98±0.07 | 0.06±0.21 | 116.19 ±252.59 | 637 012 | 0.14 | 0.82 | 0.04 |
| MOCK5 LoopSeq | Ariadne | 0.88±0.21 | 0.4±0.7 | 101.44 ±225.71 | 734 143 | 0.35 | 0.14 | 0.51 |
| MOCK20 10x | None | 0.38±0.15 | 2.28±0.62 | 186.49 ±129.32 | 503 205 | 0.97 | 0.03 | 0 |
| MOCK20 10x | Reference | 0.99±0.06 | 0.05±0.18 | 29.31 ±29.26 | 3 146 784 | 0.13 | 0.83 | 0.04 |
| MOCK20 10x | Ariadne | 0.94±0.18 | 0.22±0.64 | 13.56 ±22.69 | 6 920 196 | 0.13 | 0.05 | 0.83 |
| MOCK20 TELL-Seq | None | 0.39±0.15 | 2.37±0.59 | 160.45 ±110.51 | 623 026 | 0.98 | 0.02 | 0 |
| MOCK20 TELL-Seq | Reference | 1±0.02 | 0±0.05 | 23.83 ±27.65 | 4 067 621 | 0.01 | 0.99 | 0 |
| MOCK20 TELL-Seq | Ariadne | 0.87±0.24 | 0.47±0.89 | 21.96 ±30.74 | 4 074 321 | 0.25 | 0.04 | 0.71 |

**Figure 1.**
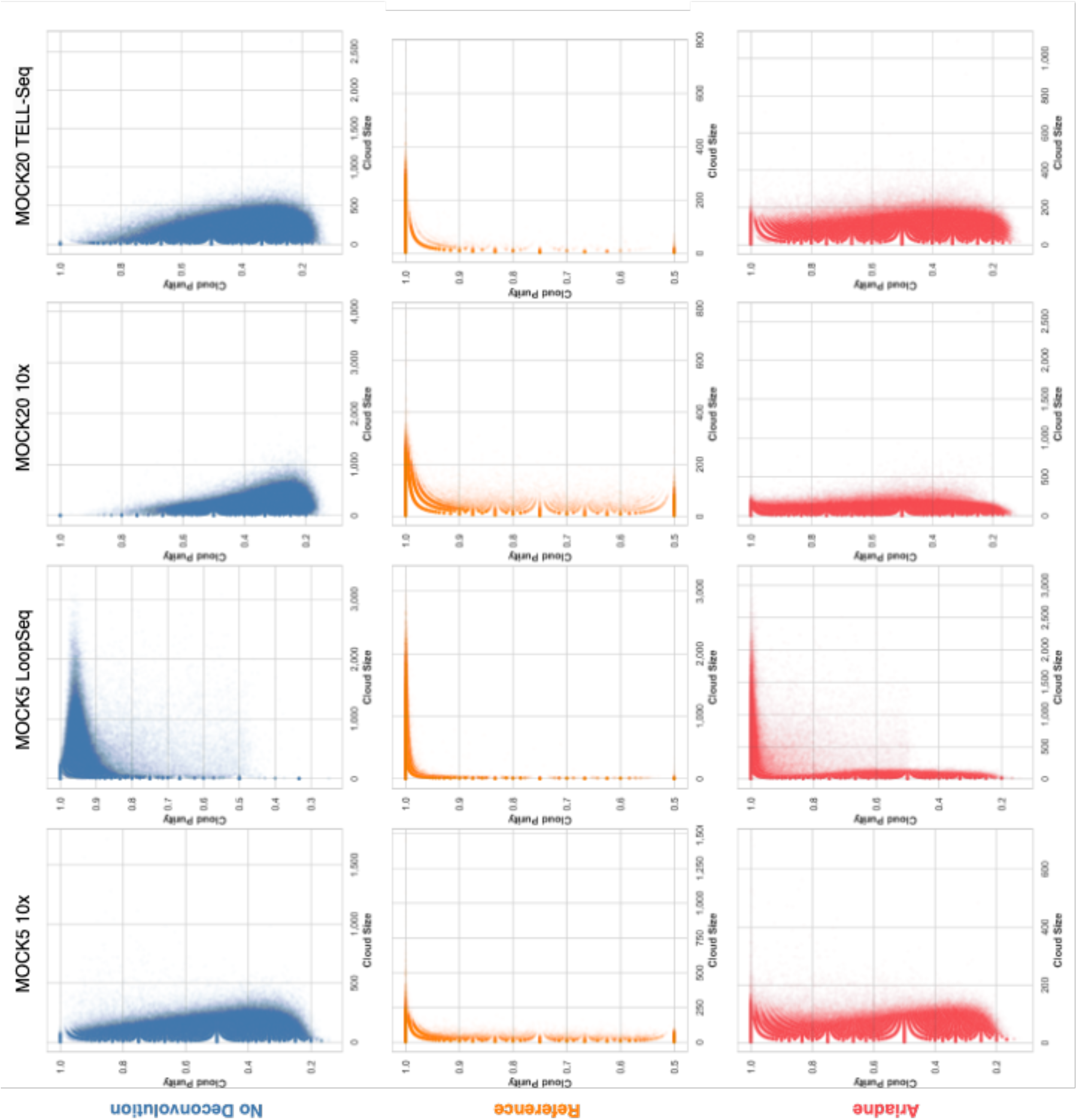
The size-weighted purity of SLR read clouds increases after applying deconvolution methods. All graphs were generated from 40,000 randomly sampled clouds from the dataset. **Top row**: No deconvolution. **Middle row**: Reference deconvolution based on read alignment to species and then grouping reads in 200 kbp regions. **Bottom row**: Ariadne deconvolution with a search distance of 5 kbp and a minimum cloud size cutoff of 6.

Ariadne at least doubles the proportion of completely deconvolved read clouds, which represent the entirety of the reads from a single inferred genomic fragment, except in the case of the MOCK5 LoopSeq dataset (Table 3 column 8). In comparison, under-deconvolved clouds are clouds comprised of reads that originate from multiple fragments, whereas over-deconvolved clouds are comprised of single-origin reads that are a subset of all of the reads from that fragment. The proportion of single-origin read clouds, the sum of complete and over-deconvolved read clouds, has increased between 2- and 37.5-fold.

While the 10x and MOCK20 TELL-Seq size-to-purity distributions are similar, the MOCK5 LoopSeq distribution peaks at P = 0.96, which explains why Ariadne deconvolution has minimal effect on read cloud quality. For the rest of the datasets, with respect to size, the relative purity of Ariadne-enhanced read clouds is significantly larger than that of the original read clouds (Figure 1 bottom row). The ideal deconvolution based on reference-mapped positions is shown in the Figure 1 middle row, where the vast majority of clouds have P = 1. This is also represented by the proportion of complete clouds, or deconvolved clouds that contain the maximal set of reads from the same original cloud that map to a single reference genome. For example, in the MOCK5 10x dataset, 91% of reference-deconvolved read clouds are complete (Table 3 row 2). Even with deconvolution, there are still large read clouds (*≥* 100 bp) in the deconvolved set, indicating that Ariadne is capable of maintaining the integrity of existing single-origin read clouds through a limited search of the assembly graph (Figure 1). Overall, applying Ariadne deconvolution to SLR datasets generates enhanced read clouds that closely resemble the size-to-purity distribution of the reference deconvolution, thereby approximating the ideal deconvolution without *a priori* information about the microbial composition of the originating data.

We have additionally demonstrated the effect of increased search distance on overall dataset quality metrics (Supp. Table 1). The number of read clouds and the average entropy increase as the search distance increases, with minimal increases in the read cloud purity along with the proportion of single-origin (complete and over-deconvolved read clouds). Given with the large increases in computational runtime, only results with the deconvolution search distance of 5 kbp are included going forward in the main text. Similar results-smaller and on average fewer-origin read clouds-can be observed with the Shannon entropy measure (Supp. Fig. 1).

### 2.4 Enhanced Read Clouds Improve Metagenomic Assembly

Since Ariadne was able to deconvolve the original SLR dataset into high-quality and non-trivial enhanced read clouds, we applied Ariadne to the full 97M-read MOCK5 10x dataset, the full 75M-read MOCK5 LoopSeq dataset, and 100M randomly sampled reads from each of the MOCK20 10x and MOCK20 TELL-Seq datasets to generate enhanced read clouds as input to cloudSPAdes [15] in metagenomics mode. The assembly quality of the resulting scaffolds with reference and Ariadne deconvolution was compared to that without prior deconvolution. cloudSPAdes generates *de novo* assemblies a wide variety of sequencing data into contigs and scaffolds, and has been benchmarked on the MOCK5 10x and MOCK20 10x datasets previously [15]. As such, we have used similar metaQUAST metrics to evaluate and compare the quality of the assemblies [30].

The largest improvements are in the overall assembly contiguity and the largest alignment, demonstrating that enhanced read clouds with increased fragment specificity generate higher-quality assemblies (Figure 2). NA50 is another measure of assembly contiguity, reporting the length of the aligned block such that using longer or equal-length contigs produces half of the bases of the assembly. *De novo* assemblies generated from reference- or Ariadne-deconvolved read clouds are significantly more contiguous than those generated from the original, non-deconvolved read clouds (Figure 2). To calculate the overall performance improvements when assembling each dataset, the assembly statistic for each species was summed and the relative difference between reference- or Ariadne-enhanced scaffolds and no-deconvolution scaffolds calculated. For NA50 and largest alignment, this means the no-deconvolution summed NA50 were subtracted from that of the reference- or Ariadne-enhanced scaffolds. For the relative misassembly rate, the number of misassembled bases was divided by the total assembly length.

**Figure 2.**
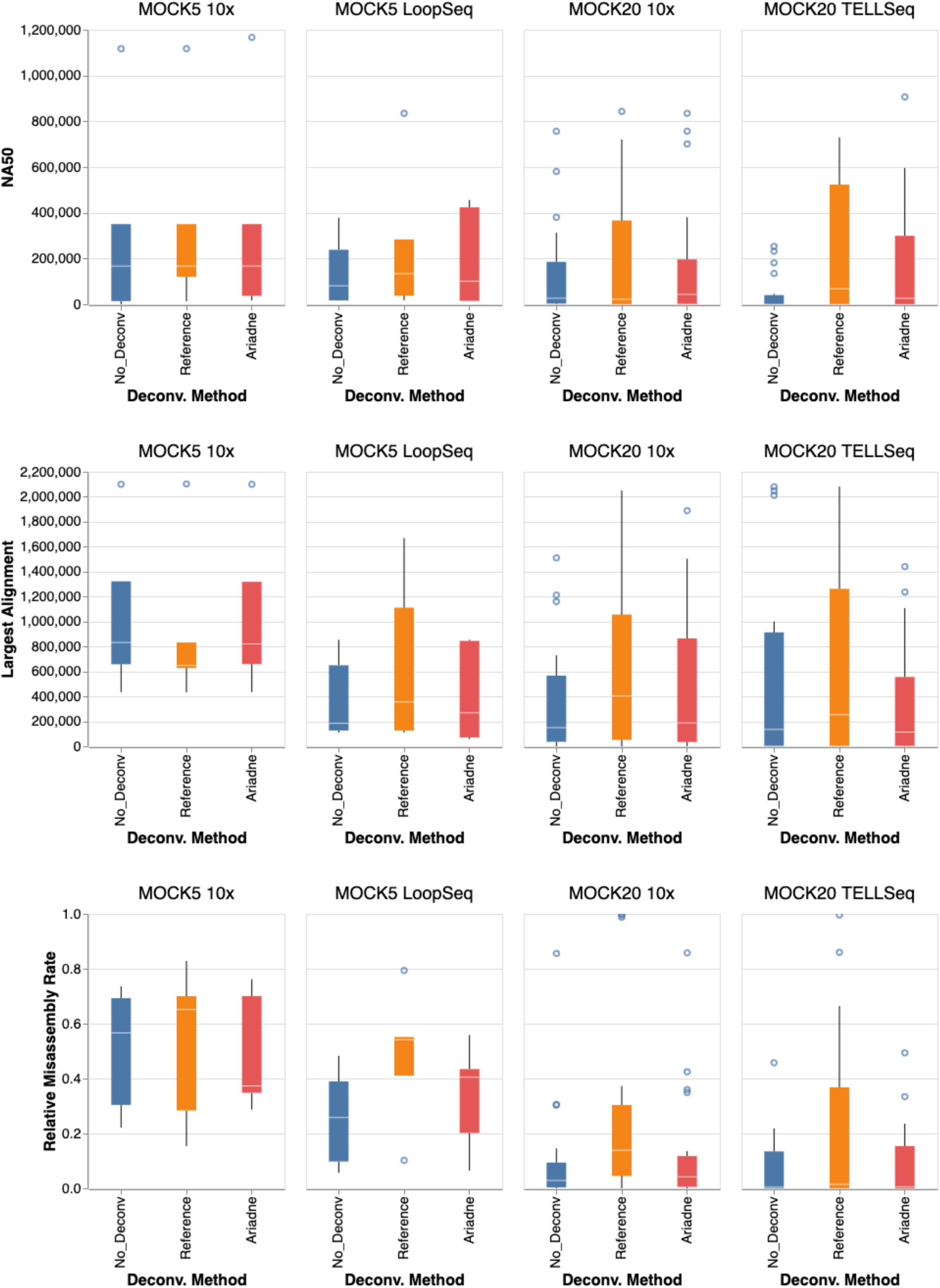
Read cloud deconvolution improves metagenomic assembly compared to raw SLR data. We compared assemblies built from raw linked reads (no deconvolution) to assemblies built from reads deconvolved using two methods: reference deconvolution, which maps reads to reference genomes, and deconvolution using Ariadne. **Top Row**: The NA50 of assemblies for each species in each sample between deconvolved reads and raw reads. Larger numbers indicate better performance. **Middle Row**: The largest alignment of assemblies for each species in each sample between deconvolved reads and raw reads. Larger numbers indicate better performance. **Bottom row**: The proportion of misassembled bases *p_miss_* is the number of bases in misassembled contigs over the total number of assembled bases.

While the differences between the deconvolution methods and no deconvolution were minimal in the MOCK5 10x dataset (Figure 2 first column), there were positive differences in NA50 and largest alignment in all other datasets. While Ariadne generally produces shorter NA50 and largest alignments than the ideal genomic deconvolution, Ariadne assemblies significantly outperform no-deconvolution assemblies in terms of contiguity statistics (Figure 2 top and middle). However, in the case of the two MOCK5 datasets, the Ariadne assembly NA50 gains were larger than the reference deconvolution assembly gains, suggesting that the most ideal read cloud partition does not always generate the most contiguous assemblies (Figure 2 top first and second panels).

In terms of misassembly rate, Ariadne assemblies largely match no-deconvolution assemblies and significantly outperform reference deconvolution. With the MOCK20 10x dataset, the third quartile of misassembly rate by reference deconvolution is 8-fold larger than that of no deconvolution, whereas Ariadne assemblies contained up to 2-fold more at maximum (Figure 2 bottom third panel). Ariadne underperforms assembly without deconvolution in terms of largest alignment and misassembly rate in MOCK20 TELL-Seq (Figure 2 bottom last panel). However, both Ariadne- and reference-enhanced assembly obtain NA50 for 4 species that the no-deconvolution assembly failed to capture, as well as some substantial differences between Ariadne and reference-based deconvolution with respect to no deconvolution (Table 4). In other datasets, there may be species that are more easily reconstructed with read cloud deconvolution that cloudSPAdes would not otherwise find long, contiguous reference subpaths for through the assembly graph. Outlier values of NA50, largest alignment, and misassembly rates were omitted from Figure 2 for visual clarity were all from the reference deconvolution scaffolds and can be found in Supp. Table 2. There were no major changes in the fraction of reference bases that were reconstructed, the total aligned length, the mean number of mismatches, and the number of contigs.

**Table 4.**
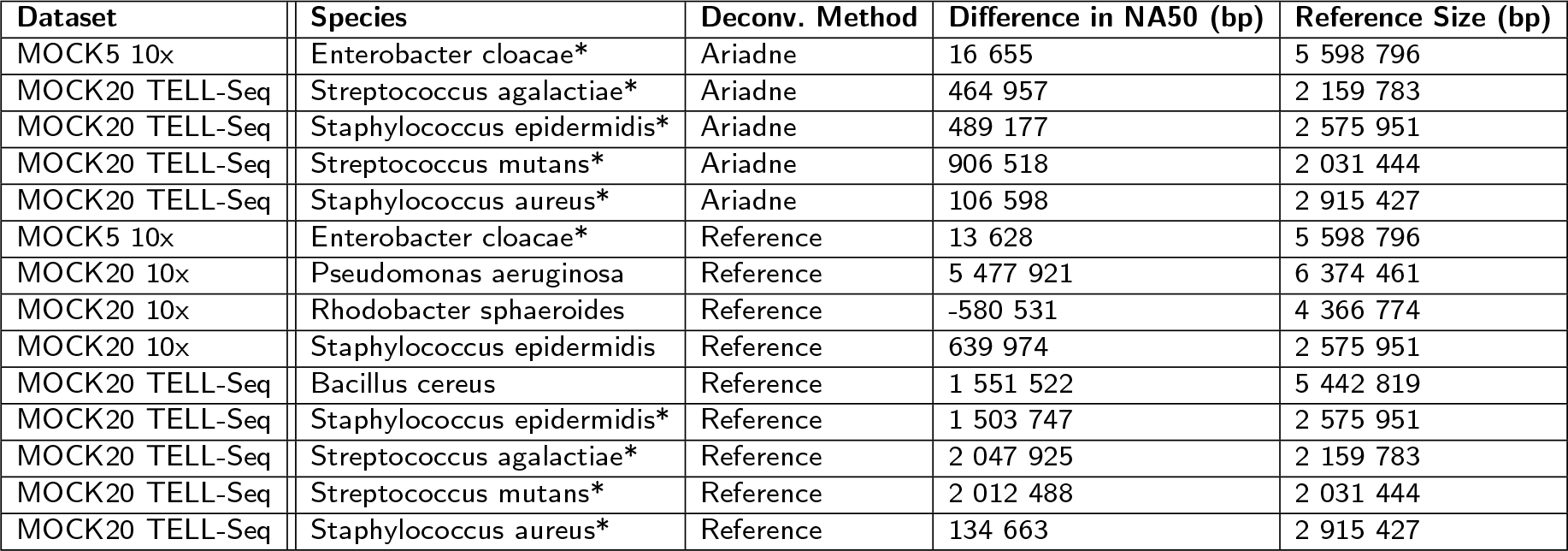
Both Ariadne and reference deconvolution increase the summed NA50 of *de novo* assembled metagenomes. The second-to-rightmost column shows the difference between the summed NA50 of assemblies obtained from deconvolved reads and the summed NA50 of non-deconvolved assembly. The asterisk (*) indicates that the assembly did not have sufficient sequence material that was alignable to the reference sequence for an NA50 to be calculated.

| Dataset | Species | Deconv. Method | Difference in NA50 (bp) | Reference Size (bp) |
| --- | --- | --- | --- | --- |
| MOCK5 10x | Enterobacter cloacae* | Ariadne | 16 655 | 5 598 796 |
| MOCK20 TELL-Seq | Streptococcus agalactiae* | Ariadne | 464 957 | 2 159 783 |
| MOCK20 TELL-Seq | Staphylococcus epidermidis* | Ariadne | 489 177 | 2 575 951 |
| MOCK20 TELL-Seq | Streptococcus mutans* | Ariadne | 906 518 | 2 031 444 |
| MOCK20 TELL-Seq | Staphylococcus aureus* | Ariadne | 106 598 | 2 915 427 |
| MOCK5 10x | Enterobacter cloacae* | Reference | 13 628 | 5 598 796 |
| MOCK20 10x | Pseudomonas aeruginosa | Reference | 5 477 921 | 6 374 461 |
| MOCK20 10x | Rhodobacter sphaeroides | Reference | -580 531 | 4 366 774 |
| MOCK20 10x | Staphylococcus epidermidis | Reference | 639 974 | 2 575 951 |
| MOCK20 TELL-Seq | Bacillus cereus | Reference | 1 551 522 | 5 442 819 |
| MOCK20 TELL-Seq | Staphylococcus epidermidis* | Reference | 1 503 747 | 2 575 951 |
| MOCK20 TELL-Seq | Streptococcus agalactiae* | Reference | 2 047 925 | 2 159 783 |
| MOCK20 TELL-Seq | Streptococcus mutans* | Reference | 2 012 488 | 2 031 444 |
| MOCK20 TELL-Seq | Staphylococcus aureus* | Reference | 134 663 | 2 915 427 |

### 2.5 Comparison of SLR Sequencing Technologies

While both comprised of 5 species, the only species common to the MOCK5 10x and MOCK5 LoopSeq datasets is *Escherichia coli*. As such, while they are comparable in terms of input community complexity, they cannot be treated as a comparison of SLR technologies with respect to *de novo* assembly. However, the two MOCK20 datasets were generated from the same 20-species mock community product from Zymo, and as such, it is possible to compare assembly quality metrics (Table 5).

**Table 5.**
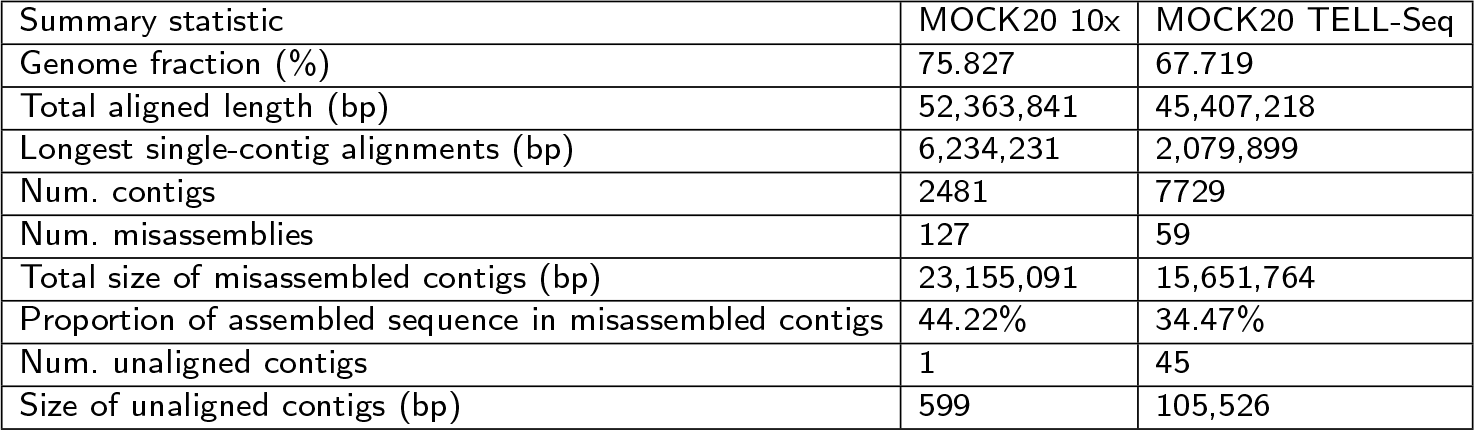
Comparison of 10x vs. TELL-Seq *de novo* assembly statistics as calculated by MetaQUAST. MOCK20 datasets were made using 10x and TELL-seq library preparation and sequencing protocols, reference-deconvolved, and *de novo* assembled using cloudSPAdes.

| Summary statistic | MOCK20 10x | MOCK20 TELL-Seq |
| --- | --- | --- |
| Genome fraction (%) | 75.827 | 67.719 |
| Total aligned length (bp) | 52,363,841 | 45,407,218 |
| Longest single-contig alignments (bp) | 6,234,231 | 2,079,899 |
| Num. contigs | 2481 | 7729 |
| Num. misassemblies | 127 | 59 |
| Total size of misassembled contigs (bp) | 23,155,091 | 15,651,764 |
| Proportion of assembled sequence in misassembled contigs | 44.22% | 34.47% |
| Num. unaligned contigs | 1 | 45 |
| Size of unaligned contigs (bp) | 599 | 105,526 |

The difference between the accurately recovered fraction of each species’ genome is small (10x is on average 8% larger). There were some species (e.g. *Bacteroides vulgatus*, approx. abundance is 0.03%) that both no-deconvolution and reference-deconvolved 10x and TELL-Seq assemblies struggled to reconstruct because of their low abundances. It is noteworthy that the Ariadne-deconvolved 10x assembly managed to reconstruct a small contig from *Schaalia odontolytica*, which both no-deconvolution and reference-deconvolved assemblies completely missed. The difference between the total alignable lengths of the assemblies is larger (13.3%), with all 10x contig-to-reference genome alignments roughly the same or significantly longer than TELL-Seq alignments. At the species level, however, there was some variability in terms of the SLR technology that reconstructed the larger alignment. This variation was probably in part due to fluctuations in coverage as well as downsampling the full sequencing read dataset for efficient comparison. For example, in the case of *Bacillus cereus*, the longest alignable 10x contig was 166,772 bp and the longest alignable TELL-Seq contig was 2,079,803 bp. However, the recovered genome fraction for both technologies was 97.03% and 98.92% respectively, indicating that while the 10x assembly of *B. cereus* was more fragmented, it was still by and large complete and gaps were likely due to coverage variability. Similar reversals can be found where the TELL-Seq assemblies of a species are more fragmented but similarly complete. Nonetheless, the 10x assembly contains far fewer contigs than the TELL-Seq assembly- 2,481 to 7,729- which is also reflected in its smaller number of larger read clouds (Table 3).

As expected with larger assemblies, the 10x assembly has significantly more misassemblies and misassembled content than the TELL-Seq data, which is probably due to the fact that more assembled bases are contained in fewer contigs. The amount of unalignable sequence content was quite small in both, and comprised < 0.2% of both assemblies.

### 2.6 Titrating the Maximum Fragment Length for Reference-based Deconvolution

To explore whether our estimate of the maximum fragment length affects assembly quality, we reference-deconvolved and assembled two datasets made from different linked-read technologies-MOCK5 LoopSeq and MOCK20 TELL-Seq-with the parameter set to 100 kbp and 400 kbp (Figure 3, Supp. Table 4, Supp. Figure 2). In both cases, the purity, entropy, cloud size and assembly summary statistics are extremely similar to the 200 kbp reference deconvolution. In fact, there seems to be an upper limit for fragment size estimates in terms of deconvolution performance. Increasing the maximum fragment length past 200 kbp slightly decreases the average purity of MOCK20 TELL-Seq read clouds (Supp. Table 4 row 6).

**Figure 3.**
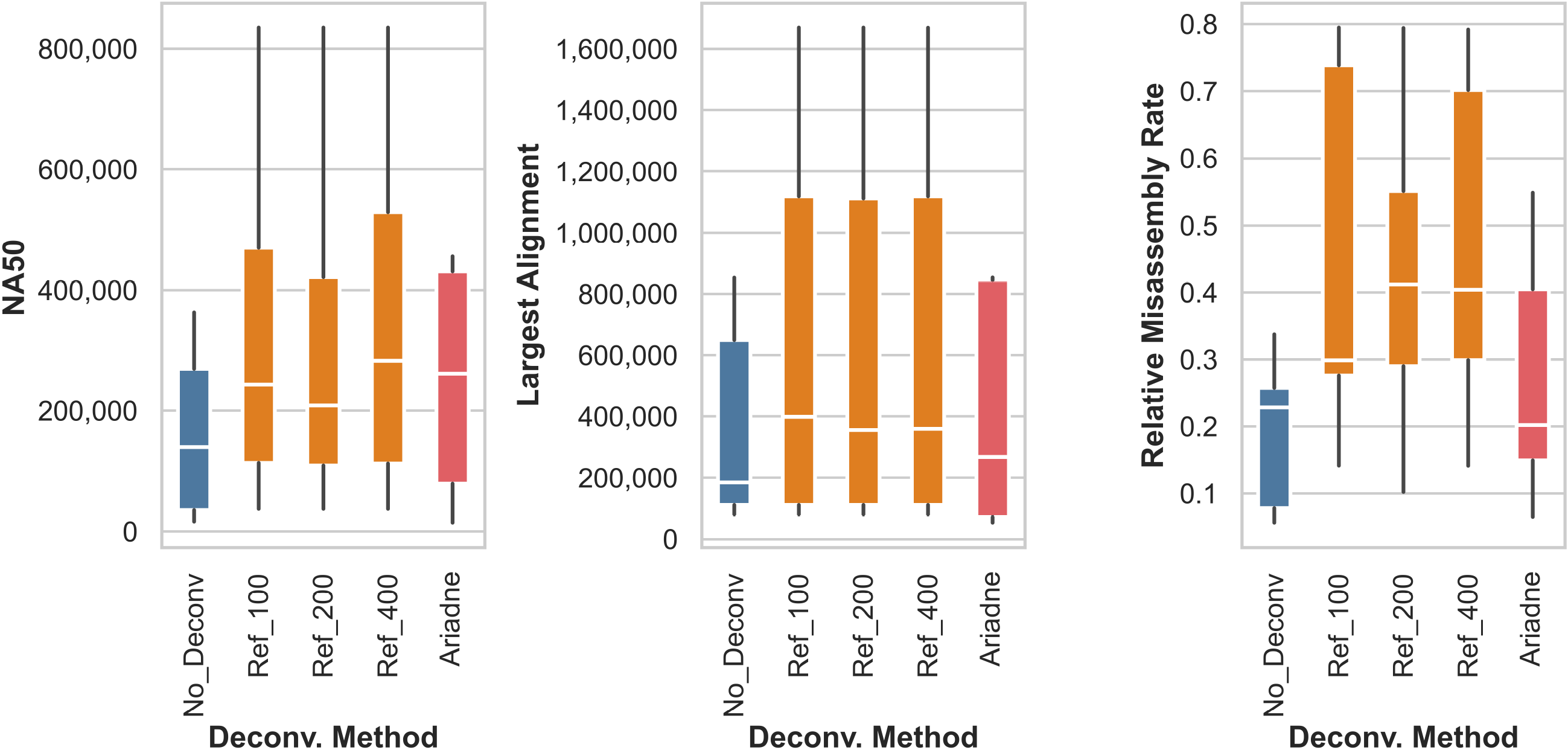
Halving and doubling the maximum fragment length does not meaningfully change the quality of de novo assembly using reference-deconvolved linked-reads. Shown here are the NA50, largest alignments, and relative misassembly rate of the MOCK5 LoopSeq reference-deconvolved assembly. As before, we compared assemblies built from raw linked reads (no deconvolution) to assemblies built from reads deconvolved using two methods: reference deconvolution with maximum fragment lengths set to 100 kbp (Ref 100), 200 kbp (Ref 200), and 400 kbp (Ref 400), and deconvolution using Ariadne.

While the median relative misassembly rate of the MOCK5 LoopSeq 100 kbp-deconvolved dataset was much lower than that of the 200 kbp-deconvolved dataset (Figure 3 rightmost panel), there was no other notable differences. It is possible that the range of values that we have explored does not generate meaningfully different enhanced cloud read compositions, and if the maximum fragment length were set lower (ex. 50 kbp) we would observe more distinctions in both the cloud properties and assembly statistics. However, the specific setting would depend on the user’s familiarity with the linked-read fragmentation protocol, while our focus was to include the maximal number of reads from a single reference sequence.

### 2.7 Barcode Deconvolution on Simulated Datasets

To evaluate the utility of barcode deconvolution on higher-complexity datasets, we simulated 100 million 10x linked reads for sets of 50 or 100 species using LRSim [31]. The species’ reference genomes were randomly selected from the United Human Gut Genome [17]. Afterwards, we conducted reference- and Ariadne-based deconvolution (search distance of 5 kbp) on the simulated datasets, then assembled the non-deconvolved, reference-deconvolved, and Ariadne-deconvolved datasets using cloudSPAdes. As previously, there is a sizable increase in the average purity of Ariadne-deconvolved clouds relative to nondeconvolved reads in the 50-species dataset (Supp. Table 4). As expected, with a larger number of species, the non-deconvolved read clouds are less pure and higher in entropy than both MOCK20 datasets, and the number of initial read clouds is approximately the same as the 10x dataset but not the TELL-Seq one. While Ariadne deconvolution does improve the overall purity and reduce the read cloud size, the differences are not as stark as with the mock microbiome datasets (Supp. Table 4). Instead of the trend observed above, where Ariadne-deconvolved read clouds are smaller than the reference-deconvolved clouds, here, they are larger instead. This may be indicative of an upper bound as to search distance-based deconvolution in terms of being able to specifically cluster sequences to species-specific clouds.

In terms of *de novo* assembly, there were minimal differences between the NA50, largest assembly, and relative misassembly rates of non-, reference-, and Ariadne-deconvolved simulated datasets (Supp. Figure 3), which was unusual given the trends in the real datasets. This could possibly be due to the average coverage of the species in the simulated datasets, which is very high-with 100 million reads to cover 160,092,218 bp and 334,517,571 bp in the 50- and 100- species datasets respectively, the depth per base is approximately 94X and 45X respectively. Simulated datasets may not be very representative of actual linked-read sequencing data, where missing genomic information due to physical or chemical wet-lab protocols are less random and less correctable through assembly. Real datasets that are not generated from mock communities may share similar missing genomic information issues as the mock communities at a larger scale. Thus, the effects of barcode deconvolution will likely be largely dependent on fragment coverage of the genome as well as the number of reads per original cloud.

### 2.8 Using Synthetic Long Reads to Improve Taxonomic Classification

Read cloud deconvolution improves short-read taxonomic classification by using, if available, the majority consensus classification of the reads in a read cloud to ‘promote’ poorly classified reads, or reclassify them at a lower-ranked taxon. Classification improvements were previously demonstrated on a test-sized dataset using Minerva-deconvolved read clouds [26], but can now be observed at the scale of full synthetic long read datasets because of Ariadne’s improvements in deconvolution runtime. The reference-deconvolved dataset is present only as a comparison; if the species composition were already known, the classification step is largely unnecessary, except in the case of strain discovery.

Whereas the non-deconvolved MOCK5 LoopSeq dataset only promoted 873 reads from root to species level, the Ariadne-deconvolved dataset promoted 20,214 reads from essentially being unclassified to the species level, which is over 22 times more than no deconvolution (Figure 4A, Table 6). Similar results can be observed with the MOCK20 TELL-Seq dataset (Figure 4B, Supp. Table 6). The difference between the number of reads that the Ariadne-deconvolved vs. nondeconvolved datasets are able to promote decreases as we consider lower initial ranks such as order, family, etc., despite the absolute number of reads promoted using linkage information increasing with or without Ariadne’s assistance (compare rows from top to bottom in Table 6, Supp. Table 6).

**Figure 4.**
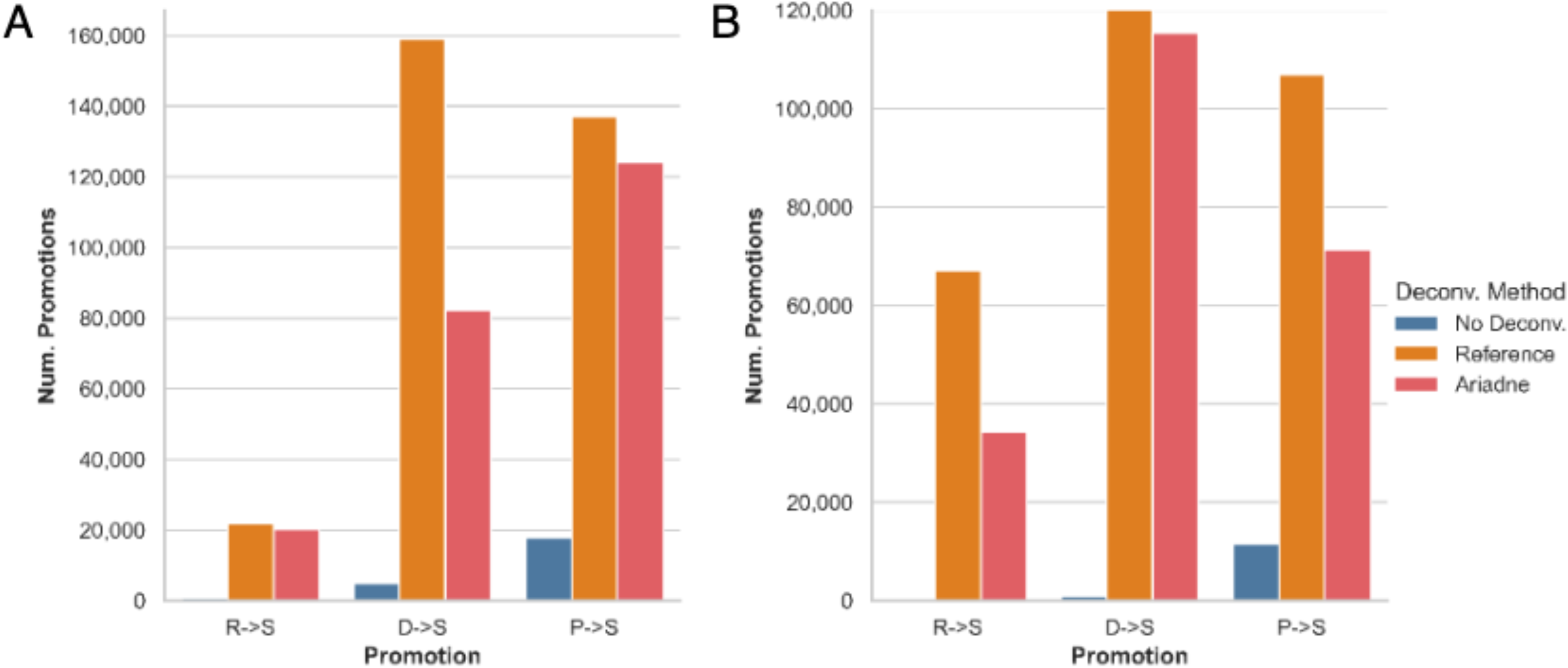
Read cloud deconvolution improves the specificity of short-read taxonomic classification, especially from high ranks such as root (R), kingdom/domain (D), and phylum (P) to low ranks such as species (S). A) MOCK5 LoopSeq. B) MOCK20 TELL-Seq, for which the y-axis has been truncated at 120,000 promoted reads for display purposes. There were 683,635 domain-to-species promotions with reference-based deconvolution.

**Table 6.** Read cloud deconvolution specifically promotes reads to low taxonomic ranks such as genus and species in the MOCK5 LoopSeq dataset. The column ‘Promotion’ indicates the promotion of a paired read i from initial rank X to rank Y as ‘X*→*Y’ using deconvolved read cloud information. The abbreviations are as follows: Root (R), kingdom/domain (D), phylum (P), class (C), order (O), family (F), genus (G), species (S). The columns ‘Prop. Improvement’ are calculated by taking the difference between the number of reads promoted using the enhanced read clouds and the number of reads promoted using the original read clouds, divided by the latter.

| Promotion | No Deconv. | Reference | Prop. Improvement | Ariadne | Prop. Improvement |
| --- | --- | --- | --- | --- | --- |
| R→D | 218728 | 46849 | -0.786 | 48874 | -0.777 |
| R→P | 17279 | 22947 | 0.328 | 29216 | 0.691 |
| R→C | 751 | 3132 | 3.17 | 4950 | 5.591 |
| R→O | 52 | 238 | 3.577 | 211 | 3.058 |
| R→F | 4416 | 109025 | 23.689 | 102446 | 22.199 |
| R→G | 732 | 40611 | 54.48 | 38472 | 51.557 |
| R→S | 873 | 21840 | 24.017 | 20214 | 22.155 |
| D→P | 43418 | 57680 | 0.328 | 62255 | 0.434 |
| D→C | 1432 | 12935 | 8.033 | 6045 | 3.221 |
| D→O | 314 | 1268 | 3.038 | 693 | 1.207 |
| D→F | 8729 | 257083 | 28.452 | 185629 | 20.266 |
| D→G | 661 | 35883 | 53.286 | 21612 | 31.696 |
| D→S | 5044 | 159161 | 30.555 | 82228 | 15.302 |
| P→C | 2919 | 5012 | 0.717 | 5263 | 0.803 |
| P→O | 173 | 695 | 3.017 | 246 | 0.422 |
| P→F | 4250 | 58696 | 12.811 | 52537 | 11.362 |
| P→G | 1477 | 27221 | 17.43 | 21011 | 13.225 |
| P→S | 17901 | 137048 | 6.656 | 123977 | 5.926 |
| C→O | 1433 | 1390 | -0.03 | 642 | -0.552 |
| C→F | 42320 | 82915 | 0.959 | 81462 | 0.925 |
| C→G | 10177 | 28636 | 1.814 | 25441 | 1.5 |
| C→S | 362453 | 376538 | 0.039 | 379622 | 0.047 |
| O→F | 374749 | 374638 | -0.0 | 372959 | -0.005 |
| O→G | 62752 | 60838 | -0.031 | 60382 | -0.038 |
| O→S | 691071 | 694087 | 0.004 | 699056 | 0.012 |
| F→G | 1096859 | 1092951 | -0.004 | 1094013 | -0.003 |
| F→S | 1354793 | 1360920 | 0.005 | 1365901 | 0.008 |
| G→S | 1957897 | 1962799 | 0.003 | 1966810 | 0.005 |

Ariadne deconvolution creates smaller purer clouds (Figure 1, bottom row), which are more likely to have originated from a constrained genomic region of a single species. Since the taxonomic classifications of Ariadne-deconvolved read clouds are more constrained than large mixed-origin non-deconvolved read clouds, poorly classified reads are much more likely to be promoted to a lower consensus rank, even if there is disagreement between the reads at the species level (Table 6 rows where the promoted-to rank is not *S*, or species, Supp. Table 6). There are some cases where the non-deconvolved dataset promoted more reads from rank *i* to rank *j*, where *j* is lower in rank than *i* (for example, R→D and F→G in tab6 and Supp. Table 6). In these cases, the reads that would have been promoted to rank *j* were promoted to lower ranks instead. For example, compare the number of reads promoted from family to genus (F→G, non-deconvolved 1,096,859 vs. Ariadne 1,094,013) in tab6 and Supp. Table 6 to the number promoted to species (F→S, non-deconvolved 1,354,793 vs. Ariadne 1,365,901). For a species-specific example of this trend, see Supp. Table 7 to observe the tendency of enhanced read clouds to promote reads to lower ranks in *Rhodobacter sphaeroides*.

**Table 7.** Read cloud deconvolution specifically promotes reads to low taxonomic ranks such as genus and species in the MOCK5 LoopSeq dataset. The column ‘Promotion’ indicates the promotion of a paired read i from initial rank X to rank Y as ‘X→Y’ using deconvolved read cloud information. The abbreviations are as follows: Root (R), kingdom/domain (D), phylum (P), class (C), order (O), family (F), genus (G), species (S). The columns ‘Prop. Improvement’ are calculated by taking the difference between the number of reads promoted using the enhanced read clouds and the number of reads promoted using the original read clouds, divided by the latter.

| Dataset | Search Distance (bp) | Time (HH:MM) | Memory (GB) | Threads |
| --- | --- | --- | --- | --- |
| MOCK5 10x | Assembly | 10:16 | 103 | 20 |
| MOCK5 10x | 5000 | 00:48 | 56 | 20 |
| MOCK5 10x | 10000 | 00:53 | 56 | 20 |
| MOCK5 10x | 15000 | 01:16 | 56 | 20 |
| MOCK5 10x | 20000 | 00:56 | 56 | 20 |
| MOCK5 10x | Reference | 18:28 | 100 | 20 |
| MOCK5 LoopSeq | Assembly | 01:13 | 22 | 20 |
| MOCK5 LoopSeq | 5000 | 00:44 | 40 | 20 |
| MOCK5 LoopSeq | 10000 | 02:04 | 40 | 20 |
| MOCK5 LoopSeq | 15000 | 03:00 | 41 | 20 |
| MOCK5 LoopSeq | 20000 | 01:51 | 41 | 20 |
| MOCK5 LoopSeq | Reference | 03:39 | 100 | 20 |
| MOCK20 10x | Assembly | 06:39 | 58 | 20 |
| MOCK20 10x | 5000 | 00:44 | 54 | 20 |
| MOCK20 10x | 10000 | 01:09 | 54 | 20 |
| MOCK20 10x | 15000 | 01:57 | 54 | 20 |
| MOCK20 10x | 20000 | 02:38 | 54 | 20 |
| MOCK20 10x | Reference | 15:07 | 100 | 20 |
| MOCK20 TELL-Seq | Assembly | 04:09 | 57 | 20 |
| MOCK20 TELL-Seq | 5000 | 01:53 | 56 | 20 |
| MOCK20 TELL-Seq | 10000 | 03:47 | 56 | 20 |
| MOCK20 TELL-Seq | 15000 | 08:57 | 56 | 20 |
| MOCK20 TELL-Seq | 20000 | 11:44 | 57 | 20 |
| MOCK20 TELL-Seq | Reference | 14:36 | 100 | 20 |

### 2.9 Runtime and Performance

In most cases, Ariadne consumes approximately the same amount of RAM and takes less time than *de novo* assembly, with the exceptions of some search distances in combination with MOCK5 LoopSeq and MOCK20 TELL-Seq (Table 7). In general, increasing the search distance increases the runtime by a few minutes to an additional hour, likely due to the fact that more neighboring vertices are considered per read with larger search distances, and a larger amount of time is taken to find intersections between larger sets of neighboring vertices. The runtime’s dependency on assembly graph connectivity is illustrated in by difference in runtime between the MOCK20 10x and TELL-Seq datasets. While they both have the same number of barcoded and paired-end reads (100 million), the TELL-Seq deconvolution takes nearly 5 times longer to complete than the 10x deconvolution. Crucially, in all cases (except for the 15 kbp+ search distances and MOCK5 LoopSeq), *de novo* assembly (i.e.: the process of generating the assembly graph) plus Ariadne deconvolution takes less time than reference-based deconvolution.

### 2.10 EMA, Longranger and Lariat, and Minerva

EMA takes SLR UMIs into account to align linked reads to a reference sequence(s) using a latent variable model, thereby serving as an alternative to Bowtie2-based read alignment to generate gold-standard read cloud deconvolution. However, the first step of the EMA pipeline (last downloaded: May 7, 2021), ema count, which counts UMIs and partitions the original FastQ file into a number of deconvolution bins, is unable to recognize most to all of the UMIs, even when custom whitelists with the exact list of UMIs in the dataset are directly provided (Supp. Table 8). In one case, ema count failed to detect any UMIs in the MOCK5 LoopSeq dataset altogether. In the best-case scenario with the MOCK20 10x dataset, ema count recognized 199,488 barcoded reads. The core deconvolution module in the pipeline, ema align, would have been able to deconvolve at maximum 0.002% of a 94-million read dataset (Supp. Table 8). To deconvolve reads that are not recognized as barcoded, the EMA pipeline uses the same procedure as our reference deconvolution pipeline, with bwa as the read aligner instead of Bowtie2. Due to the paucity of aligned reads and the subsequent lack of deconvolution, EMA-enhanced read clouds were not featured in this analysis. For ease of use, we decided to use our own reference-based deconvolution pipeline, which automates all of the read alignment, read-subgrouping, and FastQ generation steps with a single submission command. Similar UMI recognition issues were encountered with the Longranger align pipeline and the Lariat aligner it was based on. Similar to EMA, Lariat incorporates UMI information to align linked reads to a reference sequence(s). However, the available FastQ files for all four of the datasets were not comprised of raw Illumina BCL files or the raw output of longranger mkfastq. As such, it was not possible to apply Longranger align or Lariat to the four datasets. Minerva was similarly unable to deconvolve a sufficient number of read clouds (0.002% at best with the MOCK5 LoopSeq dataset), and it too was not featured in this full analysis. However, both Ariadne and Minerva completed a deconvolution test-run of 20-million reads from the MOCK5 10x dataset. In summary, while Minerva generated nearly 100% single-species read clouds, it was only able to deconvolve 4% of the reads in total (Supp. Figures 3 and 4, Supp. Table 9). Performance-wise, the Minerva runs were 4 times as long and consumed 3 times as much memory. The results are explained in greater detail in the Supplementary Materials (pg. 8 and 9).

## 3 Discussion

Ariadne deconvolution generates enhanced read clouds that are up to 37.5-fold more likely to be single-origin, which will improve downstream applications that depend on approximating the long-range linkage information from a single species, such as taxonomic classification and *de novo* assembly. In terms of classification, Ariadne-deconvolved read clouds are by and large (with the exception of the LoopSeq dataset) much smaller and more specific in terms of read origins than non-deconvolved read clouds. This allows unclassified short reads from a cloud where other short reads have been successfully classified to be assigned to the appropriate low-ranked taxon. For the MOCK5 LoopSeq dataset, there was a 22-fold, 15-fold, and 6-fold improvement of short reads promoted from root, kingdom/domain, and phylum respectively to the appropriate species classification. Indeed, *De novo* assemblies of mock metagenome communities are significantly more contiguous with Ariadne-enhanced read clouds, without the outsized increase in misassembly rate as observed with the ideal deconvolution strategy. In terms of the dataset-specific results, increasing the number of species did not change the shape of the size-to-purity distribution greatly, reflecting the inter-technology similarities in 3^′^ UMI-to-fragment multiplicity. As such, Ariadne is capable of improving assembly results and read cloud composition across all of the SLR strategies, and unlike EMA or the Longranger pipeline, easily facilitates reanalyses of existing SLR datasets for higher-resolution *de novo* assembly and taxonomic classification without pre-existing knowledge of the originating species. As with all other assembly-based algorithms, the degree of assembly contiguity will large depend on i) the true sample composition, which determines the intrinsic genetic heterogeneity to be resolved, and ii) the amount of raw sequencing data generated, which is a function of DNA extraction efficiency (i.e.: genomic fragment size) and sequencing coverage [32]. Not only that-there seems to be significant variability between sequencing runs too even in simulation studies, as observed in with the 50- and 100-species simulated 10x linked read datasets.

Though Ariadne relies on cloudSPAdes parameters to generate the assembly graph (e.g., iterative *k*-mer sizes), the program by itself only has two: search distance and size cutoff. The maximum search distance determines the maximum path length of the Dijkstra graphs surrounding the focal read. Since each read is modelled as the center of a genomic fragment, the search distance can be thought of as the width of the fragment. As such, it should be set as the mean estimated fragment length, as determined using other means such as *a priori* knowledge of shearing duration and intensity or mapping reads with the same UMI to known reference genomes. While the estimated mean length of a metagenomic fragment according to [15] is approximately 40 kbp, the most balanced results were obtained with a significantly shorter search distance. Of the distances tried in this analysis, to balance the highest-quality assembly results, single-origin read clouds, and computational efficiency, we recommend that the user set a search distance 5 kbp or shorter to generate their own enhanced read clouds. If the user has additional information that their SLR data is comprised of larger read clouds that are nearly single-origin, such as in the MOCK5 LoopSeq dataset, then larger size cutoffs should be tried. Similarly, if the goal is to generate as much alignable genetic material as possible without as much concern for synteny or the relative spacing of genes along the genome, then larger search distances can be tried. Reference-based deconvolution, which directly approximates genomic fragments by mapping reads to the known reference sequence composition of the sample, is better for large contiguous aligned blocks. However, the high misassembly rate is detrimental to the overall integrity of the assembly and should be carefully applied even in cases where the species composition of the sample is known. For some SLR technologies, the average genomic fragment size must be estimated prior to library preparation to ensure the correct balance of DNA and reagent molarity [9, 8, 11]. This average fragment size serves as a useful upper bound of an appropriate search distance.

This study is the first to compare the performances of multiple SLR technologies on well-characterized mock microbiomes. While 10x and TELL-Seq assemblies on a 20-species community are comparable in terms of the fraction of recovered species genomes, the 10x assembly was consistently larger, more accurate, and more contiguous than the TELL-Seq assembly while only slightly more prone to misassembly. LoopSeq is an interesting case where the overall fraction of recovered species genomes was low but the overall read cloud purity was extremely high, leading to smaller but extremely contiguous (i.e.: large alignable contigs) assemblies. While 10x is a well-characterized and thoroughly validated all-around choice, it may be advantageous to use LoopSeq on microbiomes composed of a few bacteria with small genomes. As with setting search distances, we advise researchers comparing SLR methods to consider the known complexity of their samples and to choose accordingly. Across all technologies, however, a single 3^′^ UMI/barcode can be associated with short reads from 2.3 to 6.7 fragments. We arrived at this estimate by dividing the average size of the non-deconvolved read clouds by the reference-deconvolved read clouds. Even with only 5 species, the odds of a single read cloud containing reads from *≥*1 species is at least 1 *−* (0.2)^1.3^ = 0.880.

There have been other recent developments in the SLR space, intended to extract as much linkage information from barcoded reads as possible in order to maximize recovery of input genomic information. Instead of tackling the UMI deconvolution problem, Guo et al. [33] and Weng et al. [34] innovate downstream of the *de novo* assembly problem. By using SLRs in combination with graph- or *k*-mer-based methods, SLR-superscaffolder and IterCluster attempt to generate longer and higher-quality assemblies. Either of these, paired with the largely single-origin read clouds generated by Ariadne, could potentially improve the NA50 and average alignment size generated by cloudSPAdes. Hybrid sequencing and analysis strategies in the future may still take advantage of the coverage and linkage depth of SLR datasets, even when used in combination with long reads [35, 36].

## 4 Conclusion

We have developed Ariadne, a novel SLR deconvolution algorithm based on assembly graphs that addresses the 3^′^ UMI deconvolution problem for metagenomics, and enables the complete usage of the linkage information in SLR technology. Ariadne deconvolution has the largest impact when the input microbial community is large and complex, especially for taxonomic classification, which is ideal for environmental microbial samples with minimal prior characterization [20]. Further algorithmic improvements to maximize the correspondence between 3^′^ UMIs and fragments may bring about significant improvements in assembly quality, and increase its scalability to other large-scale SLR problems, such as haplotype phasing.

## 5 Methods

### 5.1 Algorithm Overview

We have developed Ariadne, an algorithm that approximately solves the UMI deconvolution problem for SLR sequencing datasets. Ariadne deconvolves read clouds by positioning each read on a de Bruijn-based assembly graph, and grouping reads within a read cloud that are located on nearby edges of the assembly graph. The grouped reads are termed enhanced read clouds. Currently, Ariadne is implemented as a module of cloudSPAdes version 3.12.0.

#### 5.1.1 The Usage of UMI Information in De Novo Assembly

The following is a summary of cloudSPAdes’ usage of UMIs to identify contigs from assembly graphs. For a more detailed description, see the Materials and Methods section of [15]. cloudSPAdes constructs and iteratively simplifies a de Bruijn assembly graph using sequencing reads and several sizes of k-mers (by default, 21, 33, and 55 bp). The goal is to recover genomic cycles through the assembly graph that correspond to whole chromosomes. Due to read error, incomplete sequencing coverage, limited sequencing depth, and biological phenomena such as repetitive regions, de novo assemblers employ a number of heuristics to find the optimal unbranching paths through the graph [13, 14, 15]. In the case of SLRs and cloudSPAdes, UMIs provide one mechanism of identifying edges that are likely part of the same genomic cycle [15]. If read *i* carrying UMI *b* aligns to edge *i*, then edge *i* is said to be associated with UMI b. The UMI similarity between two edges is the proportion of associated UMIs in common. By using UMI similarities to identify edges that likely originated from the same genomic region, cloudSPAdes connects edges within the assembly graph to approximate chromosomes.

Due to fragment-to-UMI multiplicity, barcoded metagenomic read clouds are likely comprised of reads from a few species. Since these reads are all carrying the same UMI b, the long edges associated with UMI b may be erroneously connected to form contigs with genomic material from multiple species. Chimeric contigs are hazardous for downstream analyses, since as the binning of contigs into metagenomic-assembled genomes [6, 37].

#### 5.1.2 The Assembly Graph Conception of a Fragment

Searching through the cloudSPAdes-generated assembly graph for reads in the neighboring sequence space is equivalent to approximating the sequence content of a genomic fragment given the entirety of the sequence content in the input read dataset. By necessity, reads originating from the same fragment must i) have the same 3^′^ UMI and ii) be no more further apart than the total length of the fragment. The user can specify the search distance according to *a priori* knowledge about the median size of genomic fragments generated in the first step of SLR sequencing.

This process is diagrammed in Figure 5. Reads can be mapped to a de Bruijn graph. A read (Figure 5A blue and red bars) is said to coincide with an edge of the assembly graph (Figure 5B) if the read’s sequence aligns to the edge’s sequence. The read can also be said to coincide with the vertices bordering the edge it coincides with. For example, in Figure 5C, read *i*, which consists of string ‘TGACTGC’ coincides with an edge *i* that also contains the string ‘TGACTGC’.

**Figure 5.**
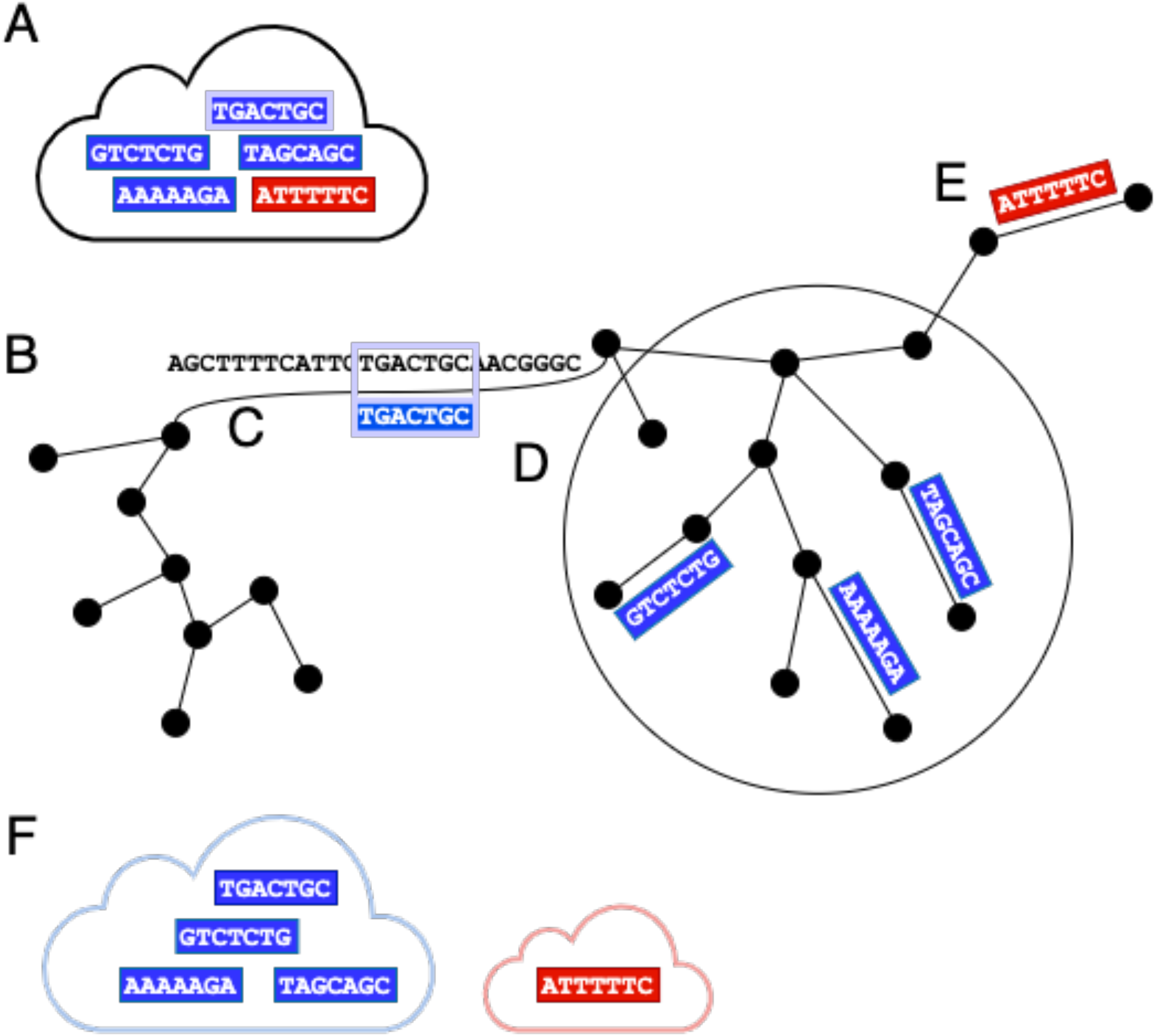
Graphical description of the Ariadne deconvolution process. **(A)** Reads with the same 3^′^ UMI are in a read cloud. Blue and red reads originate from different fragments. The **(B)** de Bruijn assembly graph is generated by cloudSPAdes, and **(C)** a focal read is mapped to one of its edges. From a read’s 3^′^-terminal vertex, **(D)** a Djikstra graph (indicated by a large black circle) is created from all edges and vertices within the maximum search distance from the 3^′^-terminal vertex. These vertices and edges (within the black circle) comprise read *i*’s search-distance-limited connected subgraph within the whole assembly graph. Reads aligning to edges in this connected subgraph are added to read *i*’s connected set. **(E)** Reads originating from different fragments likely coincide with non-included vertices. **(F)** Connected read-sets with at least one intersection (i.e.: one read in common) are output together as an enhanced read cloud.

#### 5.1.3 The Dijkstra Graph Conception of a Fragment

The Dijkstra graphs (Figure 5D) comprising the nearby assembly graph for each read represent the potential sequence space that the read-originating fragment occupies on that assembly graph. A Dijkstra graph contains the shortest paths from the source node to all vertices in the given graph. The Dijkstra graphs for a read are comprised of the vertices bordering assembly graph edges that are reachable within the maximal search distance. The maximum search distance is a user-provided parameter limiting the size of the Dijkstra graph, or the search space to be considered in the assembly graph. The maximum search distance reflects the user’s *a priori* knowledge of the size of a fragment.

### 5.2 The Ariadne Algorithm

Ariadne requires a prior step to locate reads within the (meta)genome of the sample. Where EMA requires an alignment step using the bwa read mapper to generate initial mappings for its barcoded reads [27, 38], Ariadne uses the raw assembly graph generated by any of the SPAdes family of *de novo* assemblers to identify the locations of reads relative to one another in the sample’s sequence space. Though Ariadne is intended to be a standalone tool, in the future, it will be possible to integrate Ariadne as an intermediary step in the *de novo* assembly procedure to improve the assembly graph *in situ*.

The assembly graph is first generated by applying cloudSPAdes to the raw SLR metagenomics reads as described in [39]. For a more detailed explanation of the process of forming the assembly graph, we refer the reader to [39, 40]. Subsequently, the Ariadne deconvolution algorithm is applied to the raw reads, with the cloudSPAdes-generated assembly graph supplying potential long-range linkage connections derived from the dataset.

#### 5.2.1 Step 1: Extract the assembly graph from a cloudSPAdes run

A simple representation of an assembly graph is depicted in (Figure 5B). In the cloudSPAdes assembly procedure, the assembly graph is obtained from the raw assembly graph by condensing each non-branching path into a single edge and closing gaps, removing loops, bulges, and redundant contigs. In the process of constructing the assembly graph, cloudSPAdes also maps each read to the assembly graph, generating the mapping path of a read, or the edges in the assembly graph either partially or fully spanned by the read (Figure 5C read i). The mapping path *P_i_*, or finite walk, of read *i* is comprised of the set of edges *e* from graph *G* that read *i* covers (Equation 1).

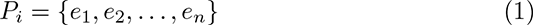

The mapping path can equivalently be described as a vertex sequence *V_i_*, which is composed of the set of vertices that border each of the edges in *P_i_* (Equation 2).

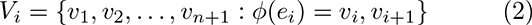

Importantly, the vertices and edges in the assembly graph are unique numeric indices replacing their sequence content. Instead of having to compare the sequences comprising the assembly graph edges, reads sharing edges can be identified by the indices in their mapping paths. As such, for the purposes of UMI deconvolution, reads do not need to be re-mapped along the assembly graph. Instead, the index-based mapping paths and vertex sequences of each read are used to locate the read. This represents a considerable speed-up over the Minerva procedure, which relies on hashing string-based sequence comparisons between reads and read clouds. Steps 2 and 3 are trivially parallelized such that the deconvolution procedure processes as many read clouds as there are threads available.

#### 5.2.2 Step 2: Generating connected read-sets for each read i

If the number of reads in the read cloud is greater than the user-set size cutoff, the read cloud (i.e.: tagged with the same 3^′^ UMI) is loaded into memory (Figure 5A). For each read *i*, the following steps are conducted to identify other reads with the same 3^′^ UMI that potentially originated from the same fragment.

##### Step 2A: Generating forward and reverse Dijkstra graphs

In Figure 5, read *i* aligns to an edge in the de Bruijn assembly graph. Assuming that read *i* is oriented in the 5*^′^ →* 3^′^ direction, the 3^′^-terminal vertex is the 3^′^-most sequence that read *i* is contiguous with. By finding edges reachable within search distance *d* of the 3^′^-terminal vertex, we will be able to all reads with the same UMI that are likely to originate from the same fragment. Reads that are oriented in the 3*^′^ →* 5^′^ direction can be reverse-complemented to apply this same procedure.

To facilitate this search, a forward Dijkstra graph (Figure 5D) is constructed starting at read *i*’s 3^′^-terminal vertex. The forward Dijkstra graph *D_f,i_* comprises the set of vertices *v_k_*, which are all vertices in the assembly graph reachable within the maximum search distance *d* from the 3^′^-terminal vertex *v_j_* (Equation 3).

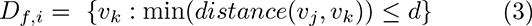

The process of constructing the Dijkstra graph, which represents the nearby sequence space of read *i*, is as follows. The tentative distances to all downstream vertices in the assembly graph are set to the maximum search distance *d*. From the 3^′^-terminal vertex, the tentative distances to these vertices are calculated from lengths of the edges connecting the vertices. This value is compared to the current distance value, and the smaller of the two is assigned as the actual distance. The vertex with the smallest actual distance from the current node is selected as the next node from which to find minimal paths, and this process is repeated. This process concludes for any path from *v_j_* when the smallest tentative distance to vertices *v_k_* is the maximum search distance.

Correspondingly, a reverse Dijkstra graph is constructed with the goal node set as read *i*’s 5’-terminal vertex, the start-vertex of the edge that the start of the read coincides with. For the reverse Dijkstra *D_r,i_*, the 5’-terminal vertex is treated as the terminal node of Dijkstra graph construction, and the set of distances in *D_r,i_* is comprised of the vertices *v_g_* such that the minimal edge-length distance between the 5’-terminal vertex and *v_g_* is less than the maximum search distance.

##### Step 2B: Identifying other reads potentially originating from the same (meta)genomic fragment

Due to long-range linkage, reads from the same genomic fragment should occur in both the nearby assembly graph and each others’ Dijkstra graphs. As such, all other reads *j* in the same read cloud are evaluated to see if they map to edges that are covered by the forward and reverse Dijkstra graphs of read i. In other words, for every other read *j* in the read cloud, if any vertex in the vertex sequence of read *j*, *V_j_* (or its reverse complement sequence) is found in read *i*’s forward or reverse Dijkstra graphs, then read *j* is added to the connected read-set of read *i* (Figure 5D).

If read j’s vertices cannot be found in the forward or reverse Dijkstra graphs of read *i*, the read is likely to have originated from a different fragment than read *i* (Figure 5E) despite being tagged with the same 3^′^ UMI, and thus no connecting component is built.

The complexity of the overall read cloud deconvolution, which consists of Dijkstra graph construction and binary searches through sets of vertices is

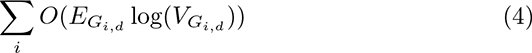

where *G* is the Djikstra subgraph induced on the overall assembly graph with read *i* as the focal read, *d* as the maximum search distance, E is the number of edges in the Djikstra subgraph *G*, and *V* is the number of vertices in the Djikstra subgraph G. The number of vertices searched for connectivity (Step 2B) depends on technical properties intrinsic to the dataset—such as the number of reads, the read coverage, error rate, number of repeats in the metagenome, and number of chromosomes—and the user-set search distance parameter. Ariadne thus deconvolves read clouds by dynamically identifying the maximal sets of connected reads within each cloud.

#### 5.2.3 Step 3: Output maximal sets of connected reads, or enhanced read clouds

Each enhanced read cloud is the set of reads from the same read cloud that are part of the same searchdistance-limited connected subgraph, as identified in step 2B. These enhanced read clouds are output as solutions to the UMI deconvolution problem (Figure 5F). Each original read cloud is either subdivided into two or more enhanced read clouds, or kept intact if all of the reads were found within the search distance *d* of each other. To avoid a preponderance of trivial-sized enhanced read clouds, if there is an enhanced read cloud generated that is composed solely of a pair of reads, the two reads are instead added to a separate set of ‘disconnected’ reads that includes read-pairs that are not connected components of any other reads or do not map to the assembly graph.

### 5.3 Selection of Search Distances

The maximum search distance was selected two quantities: i) the expected size of the physical genomic fragment that generates the sequencing reads and the ii) the likelihood of observing reads some N base-pairs (bp) away from a focal read, where N is an integer. [15] had previously estimated several dataset parameters to accurately model metagenomic fragments as paths through the assembly graph. To estimate the average fragment length, a method termed single linkage clustering was used to partition reads into clusters corresponding to alignments of likely fragments to the known reference genomes of the species comprising the sample. For example, the expected fragment size of the MOCK5 10x dataset was estimated to be 39,139 bp. For a complete analysis of the genomic fragment length estimates, the number of genomic fragments per 3^′^ UMI, and overall coverage of MOCK5 10x, see the Supplementary Materials of [15].

Setting a maximum search distance of *d* = 5 kbp seemed to model linked-read genomic fragments with an expected length of 40 kbp reasonably. To limit the occurrence of under-deconvolution, the search distance is set conservatively, as a fraction of the expected genomic fragment length. Each read *i* is used as a focal read to search the assembly graph. If read *j* truly originated from the same genomic fragment of read *i*, it is significantly more likely for read *j* to occur within *d* of read *i* than a full 40 kbp away. Several search distances smaller than the estimated fragment length—5, 10, 15, and 20 kbp—were tried for this study, and it was found that search distance *d* = 5 kbp provided the best balance of deconvolution accuracy, assembly quality, and computational efficiency (Supp. Table 1).

### 5.4 Mathematical Justification of Fragment Dissimilarity

[26] provides a short summary of the mathematical model that justifies the modelling of fragments as search-distance limited connected subgraphs within a de Bruijn assembly graph. The details of the mathematical model for drawing fragments of DNA from a SLR sequencing sample are fully described in [26], the article describing Minerva, the previous iteration of Ariadne. In brief, random fragments of genomic sequence on the scale of kilo-base-pairs from a large number of species with genomes that are many mega-basepairs in length are unlikely to be similar in terms of sequence to overlap on the assembly graph, even with k-mers as small as 22 bp. For metagenomic datasets, at least 99% of read clouds consist of fragments that do not overlap in this model [26]. This is important because it means that overlapping fragments, and thus spurious connections, are uncommon and will not hinder deconvolution in most read clouds. The algorithm is theoretically capable of uniquely deconvolving 99% of read clouds by searching the assembly graph surrounding the read for connecting reads. The only drawback of this model is that it does not account for the fact that individual fragments may have similar sequences, such as those resulting from repetitive regions or highly conserved genes.

## Appendix

### Acknowledgements

We would like to thank Indira Wu and Tuval Ben-Yehezkel at Loop Genomics for providing the MOCK5 LoopSeq dataset, as well as Yu Xia and Tom Chen at Universal Sequencing for providing the MOCK20 TELL-Seq dataset. We also thank 10x Genomics and Stephen Williams for generating the original MOCK5 and MOOCK20 10x data for us.

### Funding

This work was supported by the National Institute of General Medical Sciences (NIGMS) of the National Institutes of Health (NIH), Maximizing Investigators’ Research Award (MIRA) R35GM138152 to I.H. The content is solely the responsibility of the authors and does not necessarily represent the official vi ews of the National In stitutes of He alth. L. M., D. M. and D.C.D. were also supported by the Tri-Institutional Training Program in Computational Biology and Medicine (CBM) funded by the NIH grant 1T32GM083937. L.M. was additionally supported by the Natural Sciences and Engineering Research Council of Canada (NSERC) Postgraduate Scholarship-Doctoral (PGS D). W.N.B. was supported by the Tri-Institutional Computational Biology Summer Program.

### Abbreviations

Not applicable.

### Availability of data and materials

5.1 Data

The MOCK5 and MOCK20 10x datasets originally used in [15] are available at [cloudSPAdes.tar.gz](https://s3.us-east-2.amazonaws.com/readclouds/cloudspades_data.tar.gz) and will be submitted to SRA as well. The MOCK5 LoopSeq and MOCK20 TELL-Seq datasets generated in this study have been submitted to the NCBI BioProject database under the accession number PRJNA728470.

5.2 Code

Project name: Ariadne

Project home page: [Github](https://github.com/lauren-mak/Ariadne) and Zenodo

Archived version: NA

Operating system(s): Linux Programming language: C++, Python

Other requirements: g++ (version 5.3.1 or higher), cmake (version 2.8.12 or higher), zlib, libbz2

License: [GNU General Public License, Version 2](https://github.com/lauren-mak/ariadne/blob/ariadne_2.0/LICENSE)

Other: Pipelines and scripts used to automate *de novo* assembly, summary statistics, and figure generation are available at https://github.com/lauren-mak/ariadne-assembly-support.

### Ethics approval and consent to participate

Not applicable.

### Competing interests

The authors declare that they have no competing interests.

### Consent for publication

Not applicable.

### Authors’ contributions

L.M. implemented Ariadne, and conducted all benchmarking tests. W.N.B. help with an early prototype of Ariadne. S.M. simulated the 50- and 100-species 10x linked-read datasets. N.B. assisted with analyses. D.M., D.C.D., and I.H. conceived of the initial idea, D.M. and D.C.D. provided direction and datasets to the study. All authors contributed to the writing of the manuscript. I.H. supervised the study.

### Authors’ information

Not applicable.

### Additional Files

Additional file 1 — Supplementary Materials

Supplementary tables and figures with an extended comparison of Ariadne and Minerva on a subset of the full MOCK5 10x dataset demonstrating Ariadne’s improvements in deconvolution.

## Supplementary Materials

**Supplementary Figure 1:**
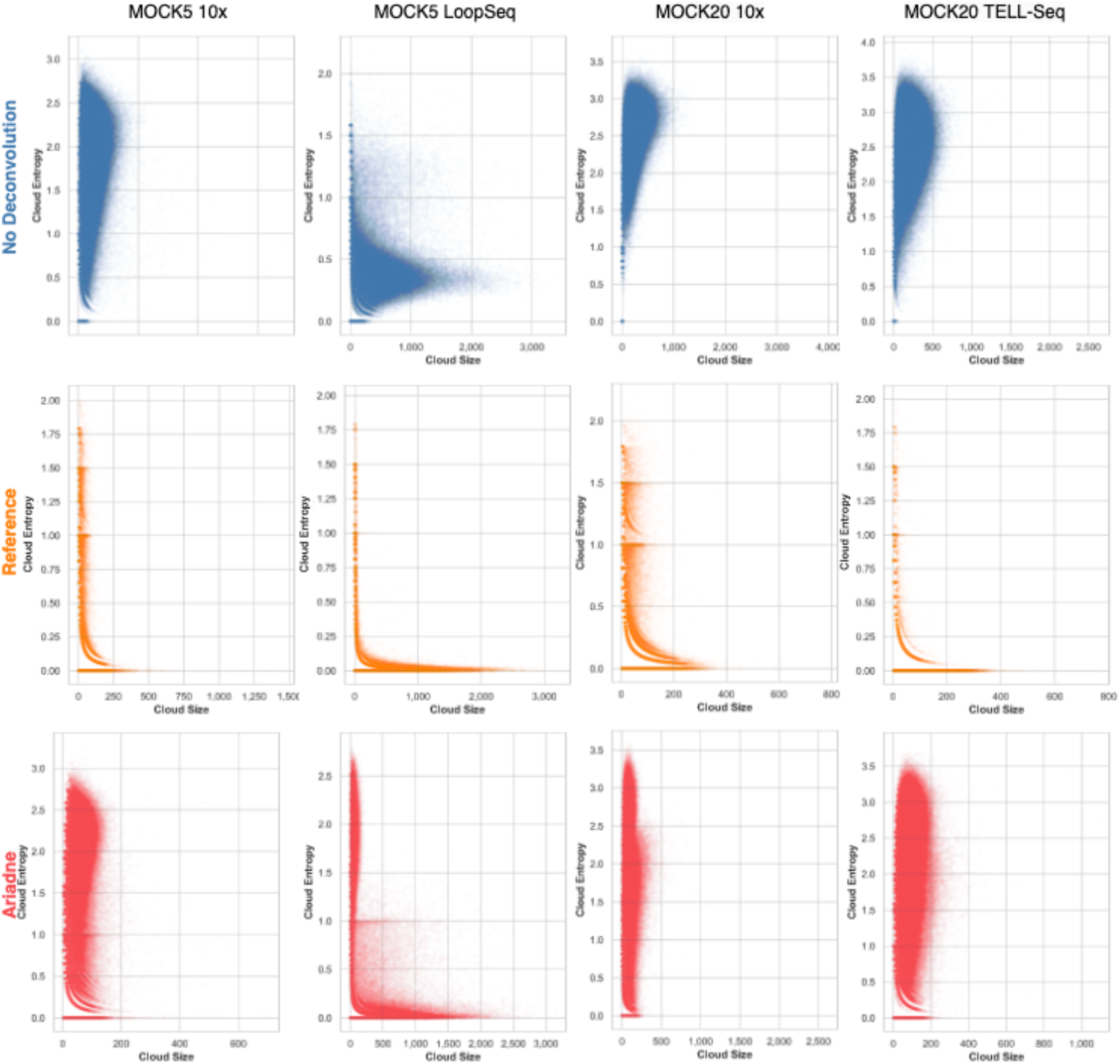
Ariadne and reference deconvolution greatly reduced large, high-entropy read clouds and create a large population of read clouds with entropy H = 0. However, the range of entropy remains the same, albeit significantly shifted towards H = 0.

**Supplementary Table 1:** Linked-read datasets of mock microbiome communities used demonstrate the utility of read cloud/barcode deconvolution prior to de novo assembly.

| Dataset Name | Num. Reads | Num. Barcoded Reads | Prop. Barcoded Reads | Species | Download | Reference Sequences | Product Sheet |
| --- | --- | --- | --- | --- | --- | --- | --- |
| MOCK5 10x | 97,491,080 | 91,101,472 | 0.93446 | Escherichia coli, Enterobacter cloacae, Micrococcus luteus, Pseudomonas fluorescens, Staphylococcus epidermidis | https://s3.us-east-2.amazonaws.com/readclouds/cloudspades_data.tar.gz | https://github.com/lauren-mak/ariadne/tree/spades_3.12.0/reference_seqs | Discontinued |
| MOCK5 LoopSeq | 75,107,814 | 75,107,814 | 1 | Escherichia coli, Porphyromonas gingivalis, Pseudomonas aeruginosa, Rhodobacter sphaeroides, Streptococcus mutans | https://www.ncbi.nlm.nih.gov/bioproject/PRJNA728470 |  | Discontinued |
| MOCK20 10x* | 100,000,000 | 94,151,528 | 0.94152 | Streptococcus mutans, Porphyromonas gingivalis, Staphylococcus epidermidis, Escherichia coli, Rhodobacter sphaeroides, Bacillus cereus, Pseudomonas aeruginosa*, Streptococcus agalactiae, Clostridium beijerinckii, Staphylococcus aureus, Acinetobacter baumannii, Neisseria meningitidis, Propionibacterium acnes, Helicobacter pylori, Lactobacillus gasseri, Bacteroides vulgatus, Deinococcus radiodurans, Actinomyces odontolyticus, Bifidobacterium adolescentis, Enterococcus faecalis | https://s3.us-east-2.amazonaws.com/readclouds/cloudspades_data.tar.gz |  | https://www.atcc.org/products/all/MSA-1003.aspx |
| MOCK20 TELLSeq* | 100,000,000 | 100,000,000 | 1 | Same as above | https://www.ncbi.nlm.nih.gov/bioproject/PRJNA728470 |  | https://www.atcc.org/products/all/MSA-1003.aspx |

**Supplementary Table 2:**
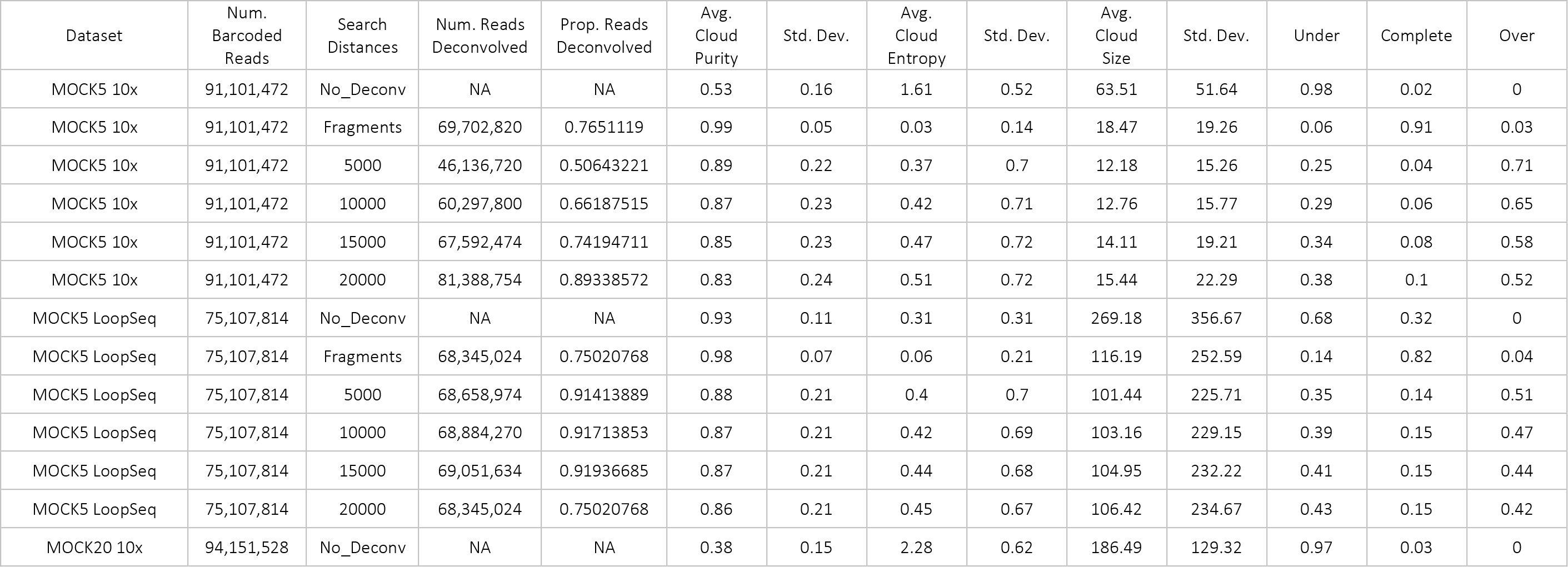
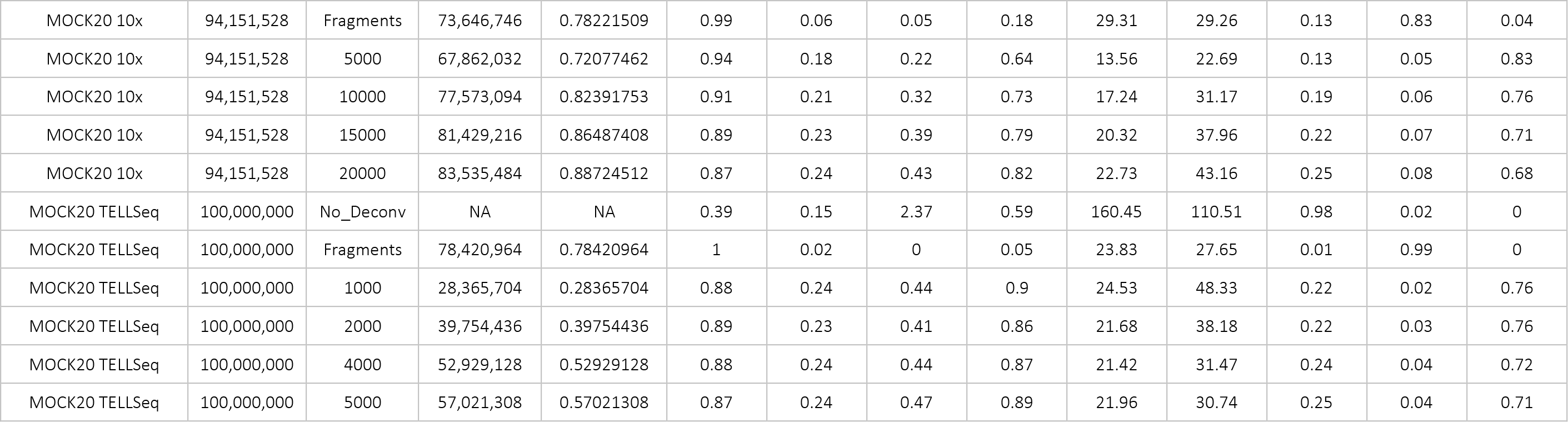
Extended version of Table 1 from the main text with additional search distances for Ariadne deconvolution. The ‘Std. Dev’ columns correspond to the parameter immediately to the left. The ‘Under, Complete, and Over’ columns refer to the proportion of total original or deconvolved read clouds that were over- or under-deconvolved, or completely and exactly comprised of all of the reads from a single inferred genomic fragment.

| Dataset | Num. Barcoded Reads | Search Distances | Num. Reads Deconvolved | Prop. Reads Deconvolved | Avg. Cloud Purity | Std. Dev. | Avg. Cloud Entropy | Std. Dev. | Avg. Cloud Size | Std. Dev. | Under | Complete | Over |
| --- | --- | --- | --- | --- | --- | --- | --- | --- | --- | --- | --- | --- | --- |
| MOCK5 10x | 91,101,472 | No_Deconv | NA | NA | 0.53 | 0.16 | 1.61 | 0.52 | 63.51 | 51.64 | 0.98 | 0.02 | 0 |
| MOCK5 10x | 91,101,472 | Fragments | 69,702,820 | 0.7651119 | 0.99 | 0.05 | 0.03 | 0.14 | 18.47 | 19.26 | 0.06 | 0.91 | 0.03 |
| MOCK5 10x | 91,101,472 | 5000 | 46,136,720 | 0.50643221 | 0.89 | 0.22 | 0.37 | 0.7 | 12.18 | 15.26 | 0.25 | 0.04 | 0.71 |
| MOCK5 10x | 91,101,472 | 10000 | 60,297,800 | 0.66187515 | 0.87 | 0.23 | 0.42 | 0.71 | 12.76 | 15.77 | 0.29 | 0.06 | 0.65 |
| MOCK5 10x | 91,101,472 | 15000 | 67,592,474 | 0.74194711 | 0.85 | 0.23 | 0.47 | 0.72 | 14.11 | 19.21 | 0.34 | 0.08 | 0.58 |
| MOCK5 10x | 91,101,472 | 20000 | 81,388,754 | 0.89338572 | 0.83 | 0.24 | 0.51 | 0.72 | 15.44 | 22.29 | 0.38 | 0.1 | 0.52 |
| MOCK5 LoopSeq | 75,107,814 | No_Deconv | NA | NA | 0.93 | 0.11 | 0.31 | 0.31 | 269.18 | 356.67 | 0.68 | 0.32 | 0 |
| MOCK5 LoopSeq | 75,107,814 | Fragments | 68,345,024 | 0.75020768 | 0.98 | 0.07 | 0.06 | 0.21 | 116.19 | 252.59 | 0.14 | 0.82 | 0.04 |
| MOCK5 LoopSeq | 75,107,814 | 5000 | 68,658,974 | 0.91413889 | 0.88 | 0.21 | 0.4 | 0.7 | 101.44 | 225.71 | 0.35 | 0.14 | 0.51 |
| MOCK5 LoopSeq | 75,107,814 | 10000 | 68,884,270 | 0.91713853 | 0.87 | 0.21 | 0.42 | 0.69 | 103.16 | 229.15 | 0.39 | 0.15 | 0.47 |
| MOCK5 LoopSeq | 75,107,814 | 15000 | 69,051,634 | 0.91936685 | 0.87 | 0.21 | 0.44 | 0.68 | 104.95 | 232.22 | 0.41 | 0.15 | 0.44 |
| MOCK5 LoopSeq | 75,107,814 | 20000 | 68,345,024 | 0.75020768 | 0.86 | 0.21 | 0.45 | 0.67 | 106.42 | 234.67 | 0.43 | 0.15 | 0.42 |
| MOCK20 10x | 94,151,528 | No_Deconv | NA | NA | 0.38 | 0.15 | 2.28 | 0.62 | 186.49 | 129.32 | 0.97 | 0.03 | 0 |
| MOCK20 10x | 94,151,528 | Fragments | 73,646,746 | 0.78221509 | 0.99 | 0.06 | 0.05 | 0.18 | 29.31 | 29.26 | 0.13 | 0.83 | 0.04 |
| MOCK20 10x | 94,151,528 | 5000 | 67,862,032 | 0.72077462 | 0.94 | 0.18 | 0.22 | 0.64 | 13.56 | 22.69 | 0.13 | 0.05 | 0.83 |
| MOCK20 10x | 94,151,528 | 10000 | 77,573,094 | 0.82391753 | 0.91 | 0.21 | 0.32 | 0.73 | 17.24 | 31.17 | 0.19 | 0.06 | 0.76 |
| MOCK20 10x | 94,151,528 | 15000 | 81,429,216 | 0.86487408 | 0.89 | 0.23 | 0.39 | 0.79 | 20.32 | 37.96 | 0.22 | 0.07 | 0.71 |
| MOCK20 10x | 94,151,528 | 20000 | 83,535,484 | 0.88724512 | 0.87 | 0.24 | 0.43 | 0.82 | 22.73 | 43.16 | 0.25 | 0.08 | 0.68 |
| MOCK20 TELLSeq | 100,000,000 | No_Deconv | NA | NA | 0.39 | 0.15 | 2.37 | 0.59 | 160.45 | 110.51 | 0.98 | 0.02 | 0 |
| MOCK20 TELLSeq | 100,000,000 | Fragments | 78,420,964 | 0.78420964 | 1 | 0.02 | 0 | 0.05 | 23.83 | 27.65 | 0.01 | 0.99 | 0 |
| MOCK20 TELLSeq | 100,000,000 | 1000 | 28,365,704 | 0.28365704 | 0.88 | 0.24 | 0.44 | 0.9 | 24.53 | 48.33 | 0.22 | 0.02 | 0.76 |
| MOCK20 TELLSeq | 100,000,000 | 2000 | 39,754,436 | 0.39754436 | 0.89 | 0.23 | 0.41 | 0.86 | 21.68 | 38.18 | 0.22 | 0.03 | 0.76 |
| MOCK20 TELLSeq | 100,000,000 | 4000 | 52,929,128 | 0.52929128 | 0.88 | 0.24 | 0.44 | 0.87 | 21.42 | 31.47 | 0.24 | 0.04 | 0.72 |
| MOCK20 TELLSeq | 100,000,000 | 5000 | 57,021,308 | 0.57021308 | 0.87 | 0.24 | 0.47 | 0.89 | 21.96 | 30.74 | 0.25 | 0.04 | 0.71 |

**Supplementary Table 3:** Extension of Table 2 in the main text. Outlier values of the relative largest alignment and relative rate of misassembled bases not represented in Figure 2 in the main text. Largest alignment is the largest alignment to the reference sequences from scaffolds obtained from the deconvolution method. The larger the relative alignment, the longer it is relative to the no-deconvolution scaffolds.

| Dataset | Species | Deconv. Method | Outlier Largest Alignment |
| --- | --- | --- | --- |
| MOCK20 10x | Escherichia coli | Reference | 2,410,209 |
| MOCK20 10x | Pseudomonas aeruginosa | Reference | 6,234,231 |
| Dataset | Species | Deconv. Method | Outlier NA50 |
| MOCK20 10x | Pseudomonas aeruginosa | Reference | 6,234,231 |
| MOCK20 10x | Streptococcus agalactiae | Reference | 2,049,091 |
| MOCK20 TELL-Seq | Bacillus cereus | Reference | 1,783,115 |
| MOCK20 TELL-Seq | Staphylococcus epidermidis | Reference | 1,503,747 |
| MOCK20 TELL-Seq | Streptococcus agalactiae | Reference | 2,047,925 |
| MOCK20 TELL-Seq | Streptococcus mutans | Reference | 2,012,488 |

**Supplementary Table 4:** Halving and doubling the maximum fragment length does not meaningfully change the quality of reference-deconvolved read clouds.

| Dataset | Max. Frag. Len. (kbp) | Avg. Purity | Avg. Entropy | Avg. Size | Num. Clouds |
| --- | --- | --- | --- | --- | --- |
| MOCK5 LoopSeq | 100 | $0.96 \pm 0.10$ | $0.08 \pm 0.34$ | $114.72 \pm 300.56$ | 638,992 |
| MOCK5 LoopSeq | 200 | $0.98 \pm 0.07$ | $0.06 \pm 0.21$ | $116.19 \pm 252.59$ | 637,012 |
| MOCK5 LoopSeq | 400 | $0.99 \pm 0.06$ | $0.06 \pm 0.08$ | $120.54 \pm 276.18$ | 636,983 |
| MOCK20 TELL-Seq | 100 | $0.99 \pm 0.04$ | $0.01 \pm 0.07$ | $23.85 \pm 32.88$ | 4,069,335 |
| MOCK20 TELL-Seq | 200 | $1 \pm 0.02$ | $0 \pm 0.05$ | $23.83 \pm 27.65$ | 4,067,621 |
| MOCK20 TELL-Seq | 400 | $0.99 \pm 0.04$ | $0.01 \pm 0.02$ | $25.83 \pm 20.62$ | 4,066,890 |

**Supplementary Figure 2:**
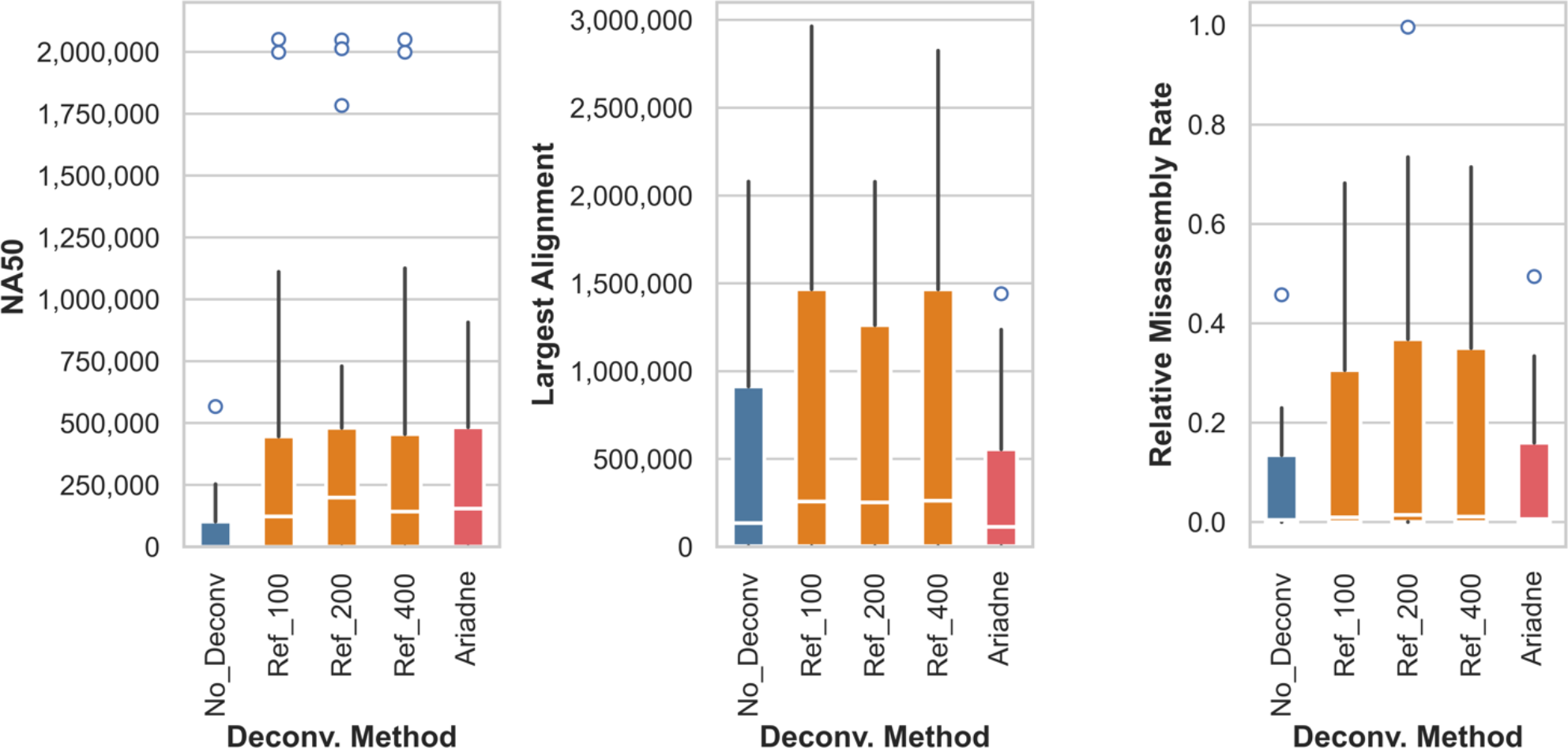
Halving and doubling the maximum fragment length does not meaningfully change the quality of de novo assembly using reference-deconvolved linked-reads. Shown here are the NA50, largest alignments, and relative misassembly rate of the MOCK20 TELL-Seq reference-deconvolved assembly.

**Supplementary Table 5:** Read cloud summary statistics for the LRSim-simulated 10x datasets of 50 and 100 species. For Ariadne deconvolution, we used a search distance of 5 kbp and for reference-based deconvolution we used 200 kbp as the maximum fragment length.

| Dataset | Deconv. Method | Avg. Cloud Purity | Std. Dev. | Avg. Cloud Entropy | Std. Dev. | Avg. Cloud Size | Std. Dev. | Num. Clouds |
| --- | --- | --- | --- | --- | --- | --- | --- | --- |
| 50-species | None | 0.29 | 0.15 | 2.39 | 0.66 | 30.37 | 15.35 | 3 290 686 |
| 50-species | Reference | 1 | 0.0029 | 0.0001 | 0.01 | 5.86 | 1.77 | 15 095 869 |
| 50-species | Ariadne | 0.45 | 0.32 | 1.85 | 1.14 | 23.57 | 16.81 | 4 237 573 |
| 100-species | None | 0.27 | 0.15 | 2.44 | 0.69 | 29.61 | 15.34 | 3 374 189 |
| 100-species | Reference | 1 | 0.001 | 0 | 0.0059 | 5.63 | 1.38 | 15 427 457 |
| 100-species | Ariadne | 0.35 | 0.26 | 2.19 | 0.97 | 26.55 | 16.24 | 3 762 776 |

**Supplementary Figure 3:**
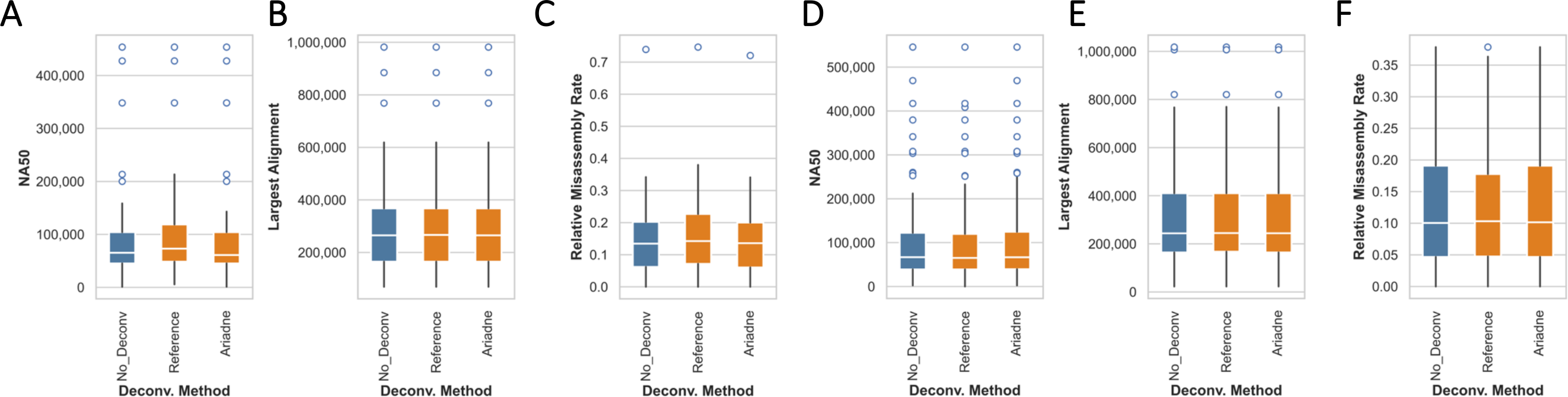
There is not much of a difference between metagenomic assembly with or without deconvolution, reference or otherwise, with linked-read datasets simulated from (A, B, C) 50 or (D, E, F) 100 species.

**Supplementary Table 6:** Read cloud deconvolution specifically promotes reads to low taxonomic ranks such as genus and species in the MOCK20 TELL-Seq dataset. The column ‘Promotion’ indicates the promotion of a paired read *i* from initial rank X to rank Y as ‘X->Y’ using deconvolved read cloud information. The abbreviations are as follows: Root (R), kingdom/domain (D), phylum (P), class (C), order (O), family (F), genus (G), species (S). The columns ‘Prop. Improvement’ are calculated by taking the difference between the number of reads promoted using the enhanced read clouds and the number of reads promoted using the original read clouds, divided by the latter. Taxon promotions where the proportion of promoted reads in the Ariadne vs. non-deconvolved datasets is greater than 10 are bolded.

| Promotion | No Deconv. | Reference | Prop. Improvement | Ariadne | Prop. Improvement |
| --- | --- | --- | --- | --- | --- |
| R->D | 330,720 | 87,665 | -0.74 | 164,568 | -0.50 |
| R->P | 207 | 380 | 0.84 | 1,236 | 4.97 |
| R->C | 87 | 633 | 6.28 | 670 | 6.70 |
| R->O | 17 | 262 | 14.41 | 142 | 7.35 |
| R->F | 180 | 155,061 | 860.45 | 123,607 | **685.71** |
| R->G | 57 | 68,966 | 1208.93 | 52,394 | **918.19** |
| R->S | 176 | 67,165 | 380.62 | 34,315 | **193.97** |
| D->P | 845 | 2,201 | 1.61 | 8,371 | 8.91 |
| D->C | 474 | 3,236 | 5.83 | 3,421 | 6.22 |
| D->O | 86 | 2,687 | 30.24 | 1,140 | **12.26** |
| D->F | 574 | 285,153 | 495.78 | 171,430 | **297.66** |
| D->G | 306 | 95,279 | 310.37 | 45,947 | **149.15** |
| D->S | 854 | 683,635 | 799.51 | 115,444 | **134.18** |
| P->C | 14,402 | 4,198 | -0.71 | 15,509 | 0.08 |
| P->O | 504 | 591 | 0.17 | 1,236 | 1.45 |
| P->F | 5,979 | 64,429 | 9.78 | 51,688 | 7.65 |
| P->G | 893 | 38,593 | 42.22 | 22,933 | **24.68** |
| P->S | 11,542 | 106,922 | 8.26 | 71,363 | 5.18 |
| C->O | 2,106 | 889 | -0.58 | 2,202 | 0.05 |
| C->F | 67,336 | 104,792 | 0.56 | 99,832 | 0.48 |
| C->G | 14,669 | 35,311 | 1.41 | 28,343 | 0.93 |
| C->S | 177,362 | 239,726 | 0.35 | 217,352 | 0.23 |
| O->F | 390,219 | 390,255 | 0.00 | 390,090 | 0.00 |
| O->G | 88,250 | 91,886 | 0.04 | 90,476 | 0.03 |
| O->S | 612,701 | 651,838 | 0.06 | 645,002 | 0.05 |
| F->G | 1,765,745 | 1,763,894 | 0.00 | 1,763,052 | 0.00 |
| F->S | 618,889 | 621,663 | 0.00 | 622,575 | 0.01 |
| G->S | 6,167,224 | 6,196,310 | 0.01 | 6,190,811 | 0.00 |

**Supplementary Table 7:** A case study of Rhodobacter (Cereibacter, using Kraken2’s database) sphaeroides taxonomic promotion. Every taxon name under ‘Promoted Rank’ is a taxon *R. sphaeroides* is classified under. The column ‘Num. Reads Difference’ is the difference between the number of reads promoted using the enhanced read clouds and the number of reads promoted using the original read clouds.

| Initial Taxon | Promoted Taxon | No Deconv. | Ariadne | Num. Reads Difference |
| --- | --- | --- | --- | --- |
| Root | Bacteria | 218,728 | 48,874 | -169,854 |
| Root | Proteobacteria | 17,279 | 29,216 | 11,937 |
| Root | Alphaproteobacteria | 5 | 5 | 0 |
| Root | Rhodobacterales | 1 | 1 | 0 |
| Root | Rhodobacteraceae | 7 | 12 | 5 |
| Root | Rhodobacter | 5 | 285 | 280 |
| Root | Rhodobacter sphaeroides | 492 | 3,587 | 3,095 |
| Bacteria | Proteobacteria | 43,404 | 62,234 | 18,830 |
| Bacteria | Alphaproteobacteria | 35 | 42 | 7 |
| Bacteria | Rhodobacterales | 29 | 84 | 55 |
| Bacteria | Rhodobacteraceae | 71 | 268 | 197 |
| Bacteria | Rhodobacter | 64 | 219 | 155 |
| Bacteria | Rhodobacter sphaeroides | 3,925 | 58,845 | 54,920 |
| Proteobacteria | Alphaproteobacteria | 51 | 219 | 168 |
| Proteobacteria | Rhodobacterales | 40 | 176 | 136 |
| Proteobacteria | Rhodobacteraceae | 128 | 525 | 397 |
| Proteobacteria | Rhodobacter | 88 | 390 | 302 |
| Proteobacteria | Rhodobacter sphaeroides | 9,041 | 101,959 | 92,918 |
| Alphaproteobacteria | Rhodobacterales | 1,232 | 486 | -746 |
| Alphaproteobacteria | Rhodobacteraceae | 2,529 | 994 | -1,535 |
| Alphaproteobacteria | Rhodobacter | 3,652 | 928 | -2,724 |
| Alphaproteobacteria | Rhodobacter sphaeroides | 354,827 | 360,909 | 6,082 |
| Rhodobacterales | Rhodobacteraceae | 2,783 | 1,026 | -1,757 |
| Rhodobacterales | Rhodobacter | 3,162 | 817 | -2,345 |
| Rhodobacterales | Rhodobacter sphaeroides | 362,506 | 368,000 | 5,494 |
| Rhodobacteraceae | Rhodobacter | 6,679 | 1,679 | -5,000 |
| Rhodobacteraceae | Rhodobacter sphaeroides | 751,157 | 758,823 | 7,666 |
| Rhodobacter | Rhodobacter sphaeroides | 694,571 | 699,793 | 5,222 |

**Supplementary Table 8:** Neither EMA nor Minerva deconvolved an appreciable part of the linked-read datasets. As such, they were omitted from the main text analysis. The specific reasons for runs that did not generate any deconvolved reads are listed in the table.

|  |  | EMA |  | Minerva |  | Reference |  |
| --- | --- | --- | --- | --- | --- | --- | --- |
| Dataset | Num. Barcoded Reads | Num. Reads Deconvolved | Prop. Reads Deconvolved | Num. Reads Deconvolved | Prop. Reads Deconvolved | Num. Reads Deconvolved | Prop. Reads Deconvolved |
| MOCK5 10x | 91,101,472 | 48,191 | 0.00049431 | Timed out after 3 days. | NA | 69,702,820 | 0.7651119 |
| MOCK5 LoopSeq | 75,107,814 | Invalid barcode<br>AAA...AA<br>whitelisted---<br>exiting! | NA | 121,238 | 0.00161419 | 54,869,732 | 0.73054625 |
| MOCK20 10x | 94,151,528 | 199,488 | 0.00199488 | 129,832 | 0.00137897 | 73,646,746 | 0.78221509 |
| MOCK20 TELLSeq | 100,000,000 | 13,553 | 0.00013553 | Timed out after 3 days. | NA | 78,420,964 | 0.78420964 |

Similar to the full 97-million read dataset, there is at least a 6.2-fold increase in the number of single-origin read clouds after Ariadne deconvolution relative to no deconvolution, regardless of the search distance. (Figure 2 top vs. bottom left). In comparison, Minerva increases the number of single-origin read clouds by 81%. Minerva is capable of deconvolving read clouds with sufficient k-mer dissimilarities between reads that originated from different fragments. However, only 4% of reads satisfies this criterion (Figures 2 bottom center vs. bottom right). This is also demonstrated by the minimal gain in average purity by the set of all read clouds that Minerva was applied to, in comparison to the subset that it was able to deconvolve (Supplementary Figure 3 and Table 6, ‘Minerva’ vs. ‘With Filter’ respectively). Thus, most of the original read clouds remain un-deconvolved with the application of Minerva. While Minerva took 15.5 hours and 163 GB of RAM, Ariadne consumed, at a maximum (search distance of 35 kbp), 4 hours and 62 GB of RAM on 20 CPUs.

**Supplementary Figure 4:**
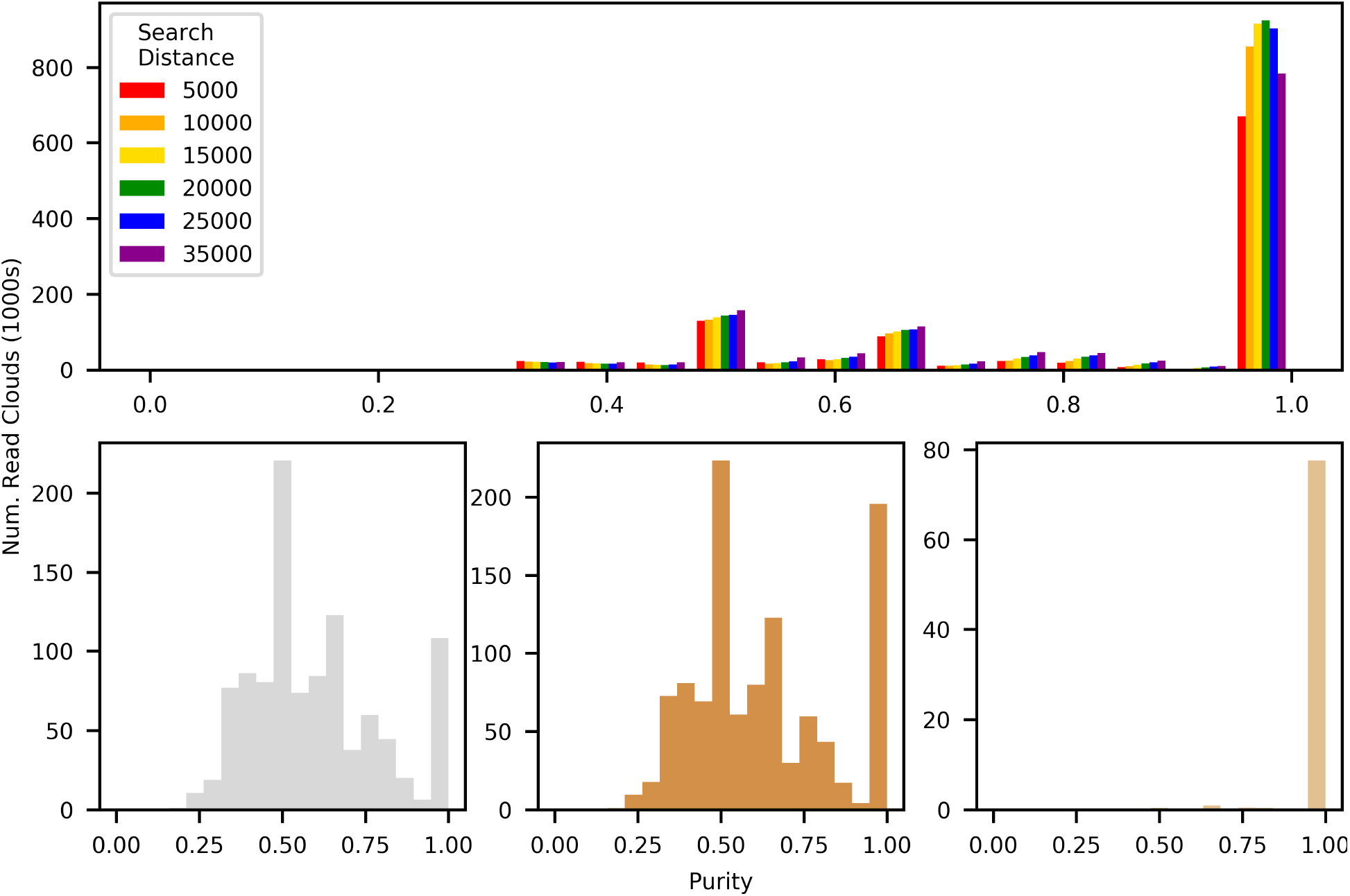
Performance comparison between Ariadne and Minerva on a 20-million read subset of the MOCK5 10x dataset. Top: Purity of Ariadne-deconvolved enhanced read clouds. Bottom left: Purity of the original read clouds. Bottom center: Purity of the entire set of read clouds after applying Minerva. This set includes read clouds that Minerva was unable to deconvolve as well as the read clouds Minerva was able to act upon. Bottom right: Purity of Minerva-deconvolved read clouds only.

**Supplementary Figure 5:**
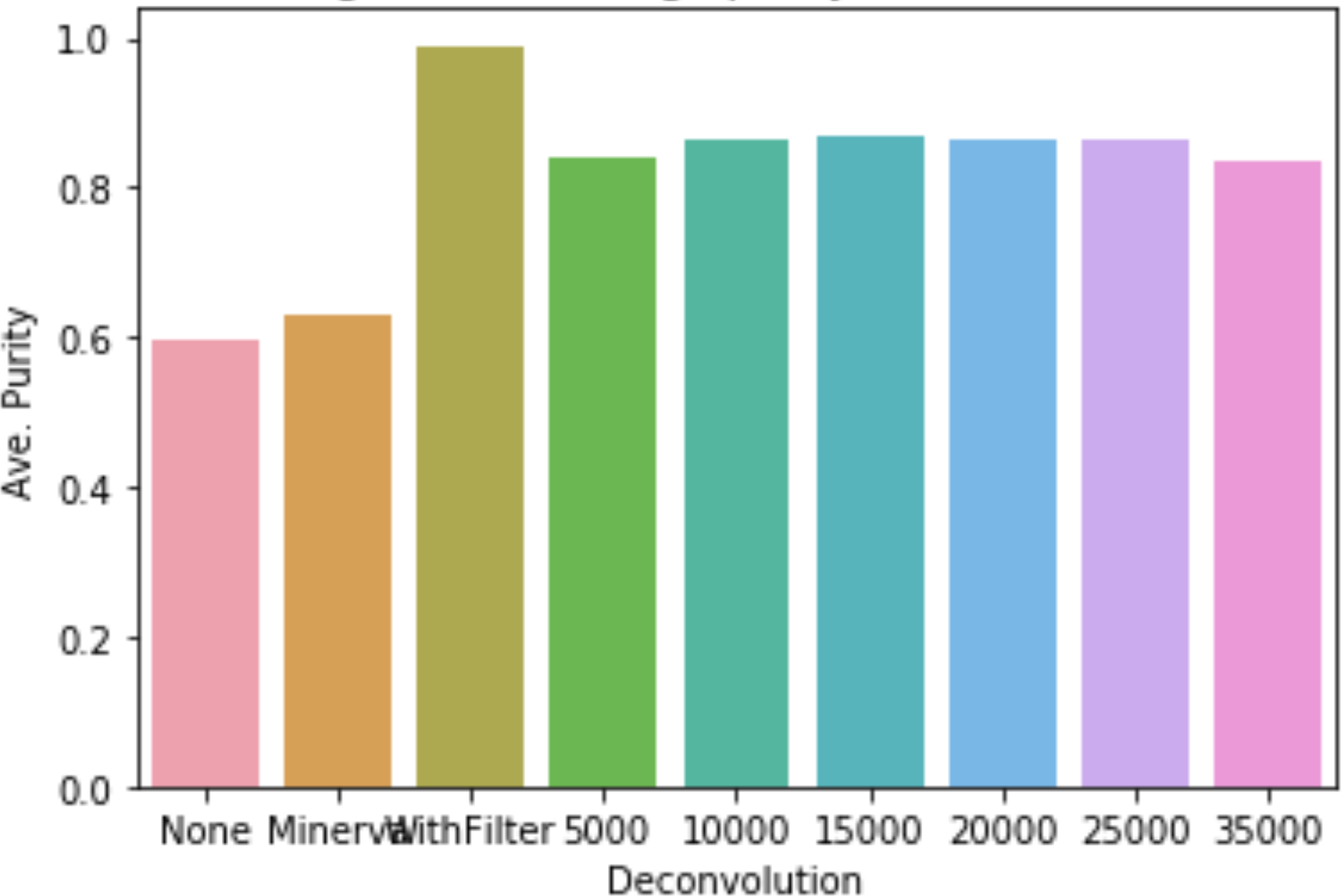
Average purity of non-deconvolved, Minerva-deconvolved with non-deconvolved, Minerva-deconvolved only (i.e.: filtered out all read clouds that could not be deconvolved by Minerva), and Ariadne-deconvolved read clouds. Ariadne deconvolution was carried out with multiple search distances (5 - 35 kbp). The same 20-million read subset of MOCK5 10x as above was used.

**Supplementary Table 9:**
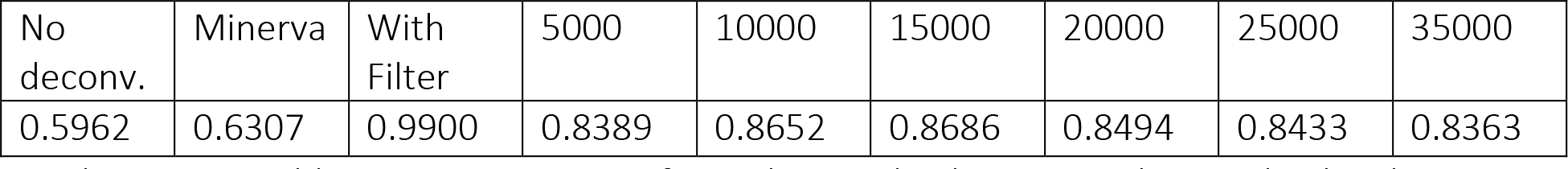
Average purity of non-deconvolved, Minerva-deconvolved with non-deconvolved, Minerva-deconvolved only, and Ariadne-deconvolved read clouds. The same 20-million read subset of MOCK5 10x as above was used.

| No deconv. | Minerva | With Filter | 5000 | 10000 | 15000 | 20000 | 25000 | 35000 |
| --- | --- | --- | --- | --- | --- | --- | --- | --- |
| 0.5962 | 0.6307 | 0.9900 | 0.8389 | 0.8652 | 0.8686 | 0.8494 | 0.8433 | 0.8363 |

